# Combinations of approved oral nucleoside analogues confer potent suppression of alphaviruses *in vitro* and *in vivo*

**DOI:** 10.1101/2025.01.24.633564

**Authors:** Sam Verwimp, Jessica Wagoner, Elijah Gabriela Arenas, Lander De Coninck, Rana Abdelnabi, Jennifer L. Hyde, Joshua T. Schiffer, Judith M. White, Jelle Matthijnssens, Johan Neyts, Stephen J. Polyak, Leen Delang

## Abstract

**Background:** Alphaviruses, including chikungunya virus (CHIKV), pose a significant global health threat, yet specific antiviral therapies remain unavailable.

**Methods:** We evaluated combinations of three oral directly acting antiviral drugs (sofosbuvir (SOF), molnupiravir (MPV), and favipiravir (FAV)), which are approved for other indications, against CHIKV, Semliki Forest virus (SFV), Sindbis virus (SINV), and Venezuelan Equine Encephalitis virus (VEEV) *in vitro* and *in vivo*. We assessed antiviral efficacy in human skin fibroblasts and liver cells, as well as in a mouse model of CHIKV-induced arthritis.

**Findings:** In human skin fibroblasts, synergistic antiviral effects were observed for combinations of MPV + SOF and FAV + SOF against CHIKV, and for FAV + SOF against SFV. In human liver cells, FAV + MPV conferred additive to synergistic activity against VEEV and SINV, while SOF synergized with FAV against SINV. In mice, MPV improved CHIKV-induced foot swelling and reduced systemic infectious virus titres. Combination treatment with MPV and SOF significantly reduced swelling and infectious virus titres compared to monotherapies of each drug. Sequencing of CHIKV RNA from joint tissue revealed that MPV caused dose- dependent increases in mutations in the CHIKV genome. Upon combination therapy of MPV with SOF, the number of mutations was significantly lower compared to monotherapy with several higher doses of MPV.

**Interpretation:** Combining these approved oral nucleoside analogues confers potent suppression of multiple alphaviruses *in vitro* and *in vivo* with enhanced control of viral genetic evolution in face of antiviral pressure. These drug combinations may ultimately lead to the development of potent combinations of pan-family alphavirus inhibitors.

**Funding:** This work was supported by a PhD fellowship granted to S.V. by the Research Foundation – Flanders (FWO) (11D5923N). L.D.C. was also supported by Research Foundation – Flanders (FWO) PhD fellowship (11L1325N). Dr. Polyak and Schiffer are partially supported by R01AI121129.

**Research in Context:** *Evidence before this study:* Alphaviruses such as chikungunya virus (CHIKV), Sindbis virus (SINV), and Venezuelan Equine Encephalitis virus (VEEV) are a major threat for global health. Alphaviruses are responsible for debilitating diseases with major public health implications, yet no antiviral drugs are currently approved for treating these virus infections. Existing treatment options are limited to supportive care and are unlikely to provide protection against future outbreaks of other alphaviruses. Previous studies have shown that oral approved nucleoside analogues such as favipiravir (FAV), molnupiravir (MPV), and sofosbuvir (SOF) have antiviral activity against certain RNA viruses, including alphaviruses. However, systematic *in vivo* evaluations of these drugs and testing of drug combinations in both *in vitro* and *in vivo* settings are limited.

*Added value of this study:* This study provides a comprehensive evaluation of combinations of FAV, MPV and SOF against multiple alphaviruses in two human cell lines and a CHIKV mouse model. We demonstrate that certain combinations of these drugs confer synergistic antiviral activity, effectively suppressing CHIKV, SFV, SINV, and VEEV replication *in vitro*. Moreover, *in vivo*, we show for the first time that MPV treatment results in reduced CHIKV-induced foot swelling and systemic virus replication. Combining MPV with SOF enhances antiviral activity in mice as compared to monotherapy. By sequencing the viral genome, we show that MPV increases the number of mutations in a dose-dependent manner. Combination therapy of MPV and SOF reduces the number of mutations compared to higher doses of MPV. These findings highlight the potential of nucleoside analogue combinations as a promising therapeutic strategy against alphavirus infections.

*Implications of all the available evidence:* The results of this study suggest that combination therapy with approved nucleoside analogues could provide an effective treatment strategy for alphavirus infections. The observed increased efficacy of drug combinations supports the potential for dose optimization to enhance efficacy while reducing toxicity and development of resistance. Future research should focus on clinical evaluation of these drug combinations, pharmacokinetic studies, and further exploration of their impact on viral evolution. Given the expanding geographical distribution of alphaviruses and the lack of available treatments, these findings provide a foundation for developing pan-alphavirus antiviral therapies that could improve patient outcomes and global outbreak preparedness.

## Introduction

Over the past 20 years, the world has experienced significant epidemics of mosquito-borne virus infections, including recurrent outbreaks of chikungunya virus (CHIKV) and other alphaviruses. Due to increased global travel and trade, global warming, and urbanization, the geographic distribution of many mosquito-borne virus infections continues to expand, including into the southeastern United States and southern Europe.^1^ Alphaviruses can cause severe diseases such as encephalitis and persistent arthritis. Over the past two decades, CHIKV has caused massive outbreaks with substantial morbidity, leading to approximately 6 million infections in Southeast Asia and the Indian Ocean Islands (2004-2006) and more than 2 million infections in the Americas (2013),^2,3^ making it the most widespread morbid alphavirus. To date, CHIKV circulation has been reported in more than 100 countries, putting 1·3 billion people at risk worldwide.^1^ Acute CHIKV infections are generally characterized by fever, skin rash, myalgia and arthralgia, which usually resolve within one week. However, CHIKV can cause lethal neurologic and cardiac symptoms. A recent study demonstrated that the case fatality rate (CFR) is 0·13%, which is higher than for other endemic arboviruses in tropical countries such as dengue virus and Zika virus.^4^ Mortality mainly affects neonates and older patients with comorbidities.^3^ Furthermore, 40 to 80% of patients suffer from chronic debilitating joint pain and swelling similar to rheumatoid arthritis, severely impacting patients’ quality of life up to months or years after the acute phase.^1,5^ Chronic CHIKV disease is estimated to result in a yearly loss of at least 158,000 disability adjusted life years and considerable economic costs.^6^

In addition to CHIKV, other members of the *Alphavirus* genus cause disease in humans, including Sindbis virus (SINV), Ross River virus (RRV), Semliki Forest virus (SFV), and the equine encephalitis viruses.^7^ SINV is widespread in Africa, Asia, Europe and Australia, while RRV is presently found in Australia, Papua New Guinea and some South Pacific islands. Both SINV and RRV cause a polyarthritic syndrome that includes fever, rash, fatigue, and painful arthritis.^8,9^ SFV is endemic to several African countries and is mostly associated with asymptomatic cases or mild febrile symptoms.^10^ Venezuelan Equine Encephalitis virus (VEEV) causes severe encephalitis in both humans and horses. Recurrent epidemics of VEEV involving substantial human infections have occurred since the turn of the 20th century.^11^ While VEEV mortality in humans is typically less than 1%, neurological symptoms occur in 14% of infected persons and are associated with long-term sequelae.^12^ Moreover, VEEV and other equine encephalitis viruses are classified as Select Agents since they are potentially weaponizable.^13^ Despite the global health threat by alphaviruses, there are currently no antiviral drugs available to treat alphavirus infections. Patients rely solely on symptomatic treatment and temporary pain relief via analgesics and nonsteroidal anti-inflammatory drugs.^14^ This highlights the clear and unmet medical need for effective anti-alphavirus therapies, particularly oral antivirals that can be easily administered. Unfortunately, there are currently no specific antiviral therapies in development for alphaviruses, with the exception of EVT894, a fully human monoclonal antibody against CHIKV, which is in phase 1 of clinical development.^15^ In addition, a live attenuated CHIKV vaccine (IXCHIQ®) was recently approved by FDA and EMA for human use, and a virus-like particle vaccine PXVX0317 is currently under review by the FDA.^16^ While vaccines are indispensable for preventing infections and reducing the disease burden, they are not sufficient to address all facets of alphavirus outbreaks. Potent and readily available antivirals will be essential to limit the severity of established infections, and possibly the duration and severity of recurrent chronic disease, and could potentially reduce transmission rates in the event of an outbreak. In this study, we evaluated oral antiviral drugs, originally approved for the treatment of other virus infections, for their ability to suppress CHIKV and related alphaviruses. By leveraging existing safety and pharmacokinetic profiles, this approach can significantly reduce the time and expense required for *de novo* drug development.^14^

Drug-based therapies that combine two or three different drugs are highly effective for the treatment of persistent chronic RNA viruses. Based on synergistic potency and prevention of drug resistance, drug combinations are the current standard of care for HCV and HIV.^17,18^

Moreover, they have shown favourable antiviral activity for RNA viruses with outbreak and pandemic potential including arenaviruses,^19^ filoviruses,^20,21^ and coronaviruses.^22–28^ Combination treatment offers multiple advantages over single dosed drugs. Combining drugs may lead to enhanced potency, particularly when there is multiplicative or synergistic activity which is most common when drugs work at different stages in the viral replication cycle. Accordingly, potent combinations may allow dose reductions leading to a decrease in drug toxicities and difficult to manage drug interactions. In addition, combination therapies dramatically lower the probability of selection for drug resistant mutants.^29^ Therefore, the use of existing approved oral drugs for emerging viruses such as CHIKV, might be improved by combining multiple drugs.

A systematic assessment of drug combinations for alphavirus infections has not yet been performed. Several studies found that a combination of the nucleoside analogue ribavirin with other drugs was more potent than single drugs in Vero,^30–33^ or U2OS cells.^34^ When FAV was combined with interferon alpha, it only slightly enhanced the antiviral effect of FAV.^35^

Herein, we demonstrate the additive to synergistic and pan-alphavirus potential of combining direct-acting RNA virus replication inhibitors. The activity of three nucleoside analogues, including favipiravir (FAV), originally approved to treat influenza virus infections in Japan;^36^ molnupiravir (MPV), which received Emergency Use Authorization in the USA for COVID;^37^ and sofosbuvir (SOF), approved in 2013 for chronic HCV infection,^38^ was assessed in human cell lines and a CHIKV infection mouse model. Our results show synergistic antiviral activity of combinations of these drugs against four different alphaviruses in two different human cell lines, highlighting their broad-spectrum antiviral potential. In mice, we demonstrated that MPV is a potent inhibitor of systemic CHIKV infections, associated with mutagenic activity. Combining MPV with SOF resulted in enhanced antiviral activity and a reduction of mutations in the CHIKV genome. Together, our results show the dual benefits of combining DAAs, paving the way for optimized therapeutic strategies against alphavirus infections.

## Materials and methods

### Cells

African green monkey kidney epithelial cells (Vero cells, ATCC CCL-81) were cultured in Minimal Essential Medium (MEM; Gibco) supplemented with 10% foetal bovine serum (FBS; HyClone), 1% L-glutamine (L-glu; Gibco), 1% sodium bicarbonate (NaHCO₃; Gibco), 1% non-essential amino acids (NEAA, Gibco) and 1% penicillin-streptomycin (Gibco). Human skin fibroblasts (ATCC CRL-2522) were maintained in MEM medium with 10% FBS, 1% L-glu, 1% NaHCO₃, 1% sodium pyruvate (Gibco), 1% NEAA and 1% penicillin-streptomycin. Human hepatocyte-derived carcinoma (Huh7) cells were cultured in MEM medium supplemented with 9% FBS, 1% NEAA, 1 mM sodium pyruvate and 1% penicillin-streptomycin. All mammalian cell cultures were incubated at 37°C, 5% CO_2_ and 95-99% relative humidity. C6/36 cells derived from *Aedes albopictus* (ATCC CRL-1660) were maintained in Leibovitz’s L-15 medium (Gibco) supplemented with 10% FBS, 1% NEAA and 1% HEPES (Gibco) and incubated at 28°C without CO_2_. Virus propagation and assays were performed using similar medium but supplemented with 2% (Vero cells, skin fibroblasts, C6/36 cells) or 5% (Huh7 cells) instead of 10% FBS.

### Viruses

The CHIKV 899 strain (GenBank FJ959103·1) was kindly provided by Prof. Drosten (University of Bonn, Germany). SFV Vietnam strain (GenBank EU350586·1) and SINV HRsp strain (GenBank J02363·1) belong to the collection of the Rega Institute of Medical Research (KU Leuven, Belgium). SINV AR86 was derived from an infectious clone provided by Dr. Mark Heise (University of North Carolina). The RRV T48 strain was obtained from the National Collection of Pathogenic Viruses (United Kingdom; 0005281v). The VEEV TC83 strain was acquired from the European Virus Archive (EVAg) or derived from an infectious clone provided by Dr. Ilya Frolov (University of Alabama). Virus stocks were produced in C6/36 cells or Vero cells and stored at -80°C. Virus titres were determined by plaque assay and end-point titrations on Vero cells. Experiments with CHIKV were conducted in the high-containment biosafety level 3 facilities of the Rega Institute of Medical Research (KU Leuven) under licenses AMV 30112018 SBB 219 2018 0892 and AMV 23102017 SBB 219 2017 0589 following institutional guidelines.

### Compounds

Molnupiravir ^39^ (MPV; EIDD-2801; EX-A3558) was purchased from Excenen Pharmatech Co., Ltd. and favipiravir ^40^ (FAV; T-705; B0084-463609 or HY-14768) was purchased from BOC Sciences or MedChem Express. Sofosbuvir ^41^ (SOF; GS-7977; HY-15005) and the cell-active metabolite of MPV, EIDD-1931 (HY-125033) were bought from MedChem Express. For *in vitro* assays, compounds were dissolved in dimethyl sulfoxide (DMSO; Sigma-Aldrich) at a concentration of 10 mM. For *in vivo* studies, MPV was dissolved in 10% PEG400 (Sigma) and 2.5% Cremophor-EL (Sigma) in water. SOF was formulated in 95% PEG400 (Sigma) and 5% Tween-80 (Sigma). FAV was dissolved in 3% NaHCO_3_ in water. For combination treatments, compounds were dissolved and administered separately to the animals. Compounds were prepared freshly every day in the morning and kept at room temperature for the second treatment of the day.

### Mice

AG129 mice were used as a model for CHIKV infection to evaluate efficacy of antiviral therapies.^42^ In-house-bred AG129 mice (deficient in IFN-α/β and IFN-γ receptors) were housed in individually ventilated isolator cages (IsoCage N Biocontainment System, Tecniplast) under standard conditions (18-23°C, 14 h:10 h light:dark cycle and 40-60% relative humidity) with cage enrichment and access to food and water ad libitum.

### Cytotoxicity and antiviral CPE reduction assay with single and combined compounds

To test the cell toxicity of compounds, Vero cells (2·5x10^4^ cells/well), skin fibroblasts (1·2x10^4^ cells/well) or Huh7 cells (5·5x10^4^) were seeded in assay medium in 96-well plates and incubated to allow adhesion overnight (37°C, 5% CO_2_). The next day, a dilution series of each single compound was prepared in assay medium on the cells. In parallel, antiviral activity was determined by adding serial dilutions of compounds to the cells, followed by infection with virus 0-2 hours after addition of compounds (MOIs are represented in Table 1). At 72 hours post-infection, cell viability and inhibition of virus-induced CPE was quantified based on metabolic activity of cells. In skin fibroblasts and Vero cells, the colorimetric MTS/PMS (Promega) method was used as viability-based read out. A 1:100 dilution of MTS reagent in colourless MEM (Gibco) was added to the cells and after incubation of 1-2h at 37°C, absorbance was measured at 490 nm (Spark, Tecan). For Huh7 cells, cell viability was determined by quantification of ATP using the CellTiter-Glo assay (Promega). A 1:2 dilution of CellTiter-Glo reagent in PBS was added, after which the plate was shaken for 2 min and incubated at room temperature for 10 min. Luminescence read out was performed on a PerkinElmer Victor X2 plate reader. Percentage of cytotoxicity and virus inhibition was calculated compared to cell control and virus control samples. The 50% cytotoxic concentration (CC_50_) and 50% effective concentration (EC_50_), the concentration of compound required to inhibit 50% of cell viability and virus-induced CPE, respectively, were calculated by logarithmic interpolation. At least three independent experiments were performed.

**Table 1.** Overview of MOIs used for antiviral assays. *MOI, multiplicity of infection*.

| Virus | Cell line | MOI |
| --- | --- | --- |
| CHIKV 899 | Vero, skin fibroblasts | 0.01 |
| SFV Vietnam | Vero, skin fibroblasts | 0.01 |
| SINV-HRsp | Vero, Huh7 | 0.01 |
| SINV-AR86 | Vero, Huh7 | 0.01 |
| RRV T48 | Vero | 0.01 |
|  | Skin fibroblasts | 0.1 |
| VEEV TC83 | Vero, skin fibroblasts, Huh7 | 0.05 |

Checkerboard assays of two drug combinations were performed as described earlier.^19^ Briefly, two compounds were tested in dose responses in 96 well plates with cells by diluting compound A across rows, while diluting compound B across columns. This checkerboard format comprises pairwise combinations of the two drugs as well as individual drugs as monotherapy. Cells were infected with virus (Table 1) and incubated until 72 hours post-infection. Percentage inhibition of virus infection was quantified based on metabolic activity of cells, as described above. Dose-response matrices with percent of virus inhibition were generated and were analysed based on the Bliss independence model of drug interactions using SynergyFinder 3·0.^43^ The degree of synergism was calculated by comparing the observed responses of drug combinations against the expected responses over the entire dose-response matrix, assuming no interaction between drugs. Bliss synergy scores higher than 10 indicate potential synergistic interactions, while scores lower than -10 indicate potential antagonism.^24^ Additionally, the Maximum Synergistic Area (MSA) score, corresponding to the maximum Bliss score calculated over a 3x3 matrix of the two compounds was represented. To evaluate the toxicity profile of these drug combinations, similar matrices of compound combinations were added to non-infected cells. Toxicity data were analysed using SynToxProfiler software.^44^ Data are represented from 2-7 independent experiments.

### Mono- and combination treatment of AG129 mice

Male AG129 mice of 6- to 10-weeks old (n=5/group) were treated with vehicle, single doses of MPV, SOF, or a combination of MPV and SOF by oral gavage. Treatment started one hour prior to CHIKV infection and was administered twice daily, until day 3 pi. MPV was administered at 10, 50, or 100 mg/kg per treatment and SOF at 40 or 80 mg/kg per treatment. For the combination treatment, MPV was dosed to 10 mg/kg and SOF to 80 mg/kg, with both compounds being separately dissolved and administered subsequently by oral gavage.

### CHIKV infection of AG129 mice

Prior to infection, AG129 mice were anesthetized by isoflurane (Iso-Vet) inhalation. Mice were then infected with 100 plaque-forming units (PFU) of CHIKV 899 strain in a total volume of 20 µL in assay medium by subcutaneous injection in the left hind footpad. Mice were monitored daily for potential altered behaviour, weight loss and disease symptoms. Additionally, swelling of the inoculated footpad was measured daily and compared to the contralateral footpad by measuring the height and width of both footpads with a digital calliper. On day 2 pi, submandibular blood was collected and centrifuged (10 000 rpm, 10 min, 4°C) to obtain serum. Mice were euthanized on day 3 pi or when showing severe CHIKV-induced disease or exceeding weight loss of more than 20% of their initial weight, by intraperitoneal injection with Dolethal (Vétoquinol). Blood was sampled by cardiac puncture and tissues, including the draining lymph node, ipsilateral and contralateral ankle joints, liver, spleen and brain, were collected to quantify viral RNA and infectious virus levels.

### CHIKV qRT-PCR

Viral RNA from serum samples was isolated using the NucleoSpin RNA virus kit (Machery-Nagel) following the manufacturer’s protocol. Tissue samples were first homogenized in Precellys tubes containing 2·8 mm zirconium oxide beads (Bertin Instruments) and TRK lysis buffer (Omega-Biotek) (lymph node, liver, spleen, brain) or assay medium (ankle joints). Homogenization was performed using an automated homogenizer (Precellys24, Bertin Instruments) with 2 cycles at 7600 rpm for 20 s, with a 20 s interval. Tissue homogenates were then centrifuged (15 000 rpm, 10 min, 4°C), after which RNA was isolated using the E.Z.N.A Total RNA kit I (Omega-Biotek) (lymph node, liver, spleen, brain) or the NucleoSpin RNA virus kit (Machery-Nagel) (ankle joints), according to the manufacturers’ protocols.

To quantify the isolated viral RNA, qRT-PCR was performed in a total reaction volume of 25 µl, containing 13·94 µl nuclease free water (Promega), 6·25 µl master mix (Eurogentec), 0·375 µl forward primer (5’-CCG ACT CAA CCA TCC TGG AT-3’; final concentration of 150 nM; IDT), 0·375 µl reverse primer (5’GGC AGA CGC AGT GGT ACT TCC T-3’; final concentration of 150 nM; IDT), 1 µl probe (5’-FAM-TCC GAC ATC ATC CTC CTT GCT GGC-TAMRA-3’; final concentration of 400 nM; IDT), 0·0625 µl reverse transcriptase (Eurogentec) and 3 µl RNA. The qRT-PCR was performed on the QuantStudio 5 Real-Time PCR system (ThermoFisher Scientific) using the following programme: 30 min at 48°C, 10 min at 95°C, 40 cycles of 15 s at 95°C and 1 min at 60°C. A standard curve was generated based on a ten-fold dilution series of reference CHIKV plasmid DNA. The limit of detection (LOD) was determined as the lowest viral load that could be detected by the qRT-PCR assay in 95% of experiments, considering the tissue weights and buffer volumes.

### End-point titrations

To quantify infectious virus levels in mouse serum and tissues, Vero cells were seeded in 96-well plates at a density of 10^4^ cells/well and incubated overnight (37°C, 5% CO_2_). The following day, tissues were first homogenized in Precellys tubes with assay medium for two cycles of 20 s at 7600 rpm, with a 20 s interval. After centrifugation (15 000 rpm, 10 min, 4°C), ten-fold serial dilutions of tissue homogenate or serum were added to the cells in triplicate. At day 3 pi, cells were microscopically scored for CHIKV-induced cytopathogenic effect (CPE) compared to uninfected cell controls. The TCID_50_/ml (tissue culture infectious dose 50%), the concentration of virus needed to infect 50% of cell cultures, was calculated using the method of Reed and Muench.^45^ The limit of quantification was determined as the lowest viral titre that could be quantified using this method, considering tissue weights and buffer volumes.

### Deep sequencing and variant analysis

Viral RNA isolated using the NucleoSpin RNA virus kit (Machery-Nagel) from contralateral ankle joints from single dose-response and combination studies in AG129 mice were used for deep sequencing. RNA extracts were synthesized into cDNA using the Protoscript II First Strand cDNA Synthesis Kit, NEBNext Ultra II Non-Directional RNA Second Strand Module, and Random Primer 6 (New England Biolabs Inc.). 25 ng of the cDNA was used for library construction with the Twist Library Preparation EF 2·0 Kit and Twist Universal Adaptor System from Twist Biosciences. The resulting libraries were pooled and additionally enriched with the Twist Comprehensive Viral Panel at 70°C for 16h to hybridize the probes. The post-capture pool was further PCR amplified for 13 cycles and final libraries were sequenced on AVITI Systems (Element Biosciences, USA) to generate 2×150Lbp paired end reads.

The resulting reads were trimmed for quality and sequencing adapters with Trimmomatic v0·39·.^46^ Reads were subsequently mapped with Bowtie2 v2·5·1 ^47^ to the genome of reference strain CHIKV 899 (FJ959103·1). Next, PCR duplicate reads were removed with samtools v1·21,^48^ before variant calling with iVar v1·4·3 ^49^ with a minimum depth of 100X. Called variants across all samples were further analysed in R. Variants present across all controls were removed to account for cell line adaptation of the virus. Differences in the mutation count, normalized for sequencing depth, were tested with one sided Wilcoxon tests. P-values were corrected for multiple testing with the Benjamini-Hochberg correction, and adjusted p-values lower than 0·05 were considered significant. Data analysis scripts are available on https://github.com/LanderDC/CHIKV_variant_analysis.

### Statihstics

All statistical details are described in the figure legends. Variant analysis was performed as described above. Other statistical analyses were performed using Graphpad Prism 10·3·1. Significant differences in foot swelling and serum virus loads were analyzed using the two-way repeated measures ANOVA with Tukey’s correction. Significant differences in virus loads in tissues were determined using the non-parametric Kruskal-Wallis with Dunn’s multiple comparisons test (comparing independent groups), applying Bonferroni correction for multiple testing. Statistical significance threshold was assessed at p values < 0·05.

### Ethics statement

Experiments with mice were performed with the approval and under the guidelines of the Ethical Committee of the University of Leuven (license P070/2019).

### Data availability

All data generated during this study are included in this article. Data analysis scripts of sequencing studies are available on https://github.com/LanderDC/CHIKV_variant_analysis. Data generated and analysed for this study, or any additional information is available from the lead contact upon request.

## Results

### Selection of DAAs for evaluating combinations against different alphaviruses

The aim of our study is to develop combinations of approved oral direct-acting antivirals (DAAs) with potent pan-alphavirus activity *in vitro* and *in vivo*. Since there are currently no specific antiviral drugs available or in late-stage clinical development for alphaviruses, we reviewed the literature to identify oral DAAs already approved for other indications or in late-stage development, with reported activity against alphaviruses. From approximately 100 drugs described as active versus alphaviruses,^14,50–54^ we identified 3 oral DAAs (polymerase inhibitors) that have *in vitro* antiviral efficacies at concentrations that are achievable in humans: MPV, FAV and SOF. EIDD-1931, the active metabolite of MPV, showed potent antiviral efficacy against several alphaviruses in mammalian cells,^55–58^ and in a VEEV mouse model.^59^ Similarly, the influenza-inhibitor FAV exhibited *in vitro* activity against a panel of alphaviruses and was effective in the treatment of CHIKV-infected mice.^60^ SOF has also demonstrated modest activity against CHIKV in human cell lines and in two different mouse models.^61^

We first confirmed the previously reported anti-alphavirus activity of MPV (active metabolite EIDD-1931) and FAV in Vero cells (Fig S1) against a broad panel of alphaviruses (CHIKV, SFV, SINV-HRsp, SINV-AR86, VEEV TC83 and RRV). MPV exerted potent dose-dependent inhibition of virus-induced CPE against all alphaviruses tested, with EC_50_ values ranging between 0·3 and 1·2 µM (Table 2). Likewise, treatment with FAV resulted in dose-dependent reduction of alphavirus infection with EC_50_ values between 13-52 µM. In contrast, no antiviral activity was observed for SOF, most likely due to lack of metabolization of this drug in Vero cells.^62^

**Table 2.** Overview of *in vitro* EC_50_ of compounds against CHIKV, SFV, SINV, RRV and VEEV; and CC_50_ values in different cell lines. *EC_50_, 50% effective concentration; CC_50_, 50% cytotoxic concentration*.

| Vero cells |  |  |  |  |  |  | Skin fibroblasts |  |  |  |  | Huh7 cells |  |  |  |
| --- | --- | --- | --- | --- | --- | --- | --- | --- | --- | --- | --- | --- | --- | --- | --- |
| CHIKV | SFV | SINV HRsp | SINV AR86 | RRV | VEEV TC83 | CC<sub>50</sub> | CHIKV | SFV | RRV | VEEV TC83 | CC<sub>50</sub> | SINV HRsp | SINV AR86 | VEEV TC83 | CC<sub>50</sub> |
| 0,7 ± 0,2 | 1,2 ± 0,4 | 0,7 ± 0,2 | 0,3 | 0,5 ± 0,2 | 1,1 ± 0,4 | > 200 | 0,9 ± 0,1 | 0,5 ± 0,2 | 1,2 ± 1,4 | 0,4 ± 0,1 | 140 ± 19 | 1,8 ± 0,3 | 0,6 ± 0,1 | 2 ± 0,5 | 22 ± 11 |
| > 100 | > 100 | > 100 | *n.d.* | > 100 | > 100 | 350 ± 14 | 15 ± 3 | 17 ± 3 | 20 ± 7 | 12 ± 3 | 280 ± 13 | 44 ± 3 | >50 | > 50 | 271 ± 95 |
| 35 ± 14 | 33 ± 6 | 52 ± 18 | 17 ± 2 | 15 ± 12 | 13 ± 0,3 | 374 ± 17 | 143 ± 5 | 382 | >200 | >200 | > 500 | 22 ± 7 | 29 ± 18 | 112,3 | >500 |

We next evaluated the antiviral dose-responses of these drugs as solo agents against the panel of alphaviruses in human cell lines which are biologically more relevant compared to Vero cells. More specifically, we used human skin fibroblasts,^63^ and human hepatoma-derived Huh7 cells,^64^ which are the actual target cells for alphavirus replication and/or pathology. In the skin fibroblasts, MPV resulted in substantial dose-dependent inhibition of all alphaviruses tested, with EC_50_ values in the sub to low single-digit micromolar range (Fig 1a, Table 2). SOF exhibited dose-dependent suppression of all alphaviruses tested in skin fibroblasts, with EC_50_ values between 12-20 µM (Fig 1b, Table 2). In contrast, FAV was only effective against CHIKV, with an EC_50_ of 143 µM (Fig 1c).

**Figure 1.**
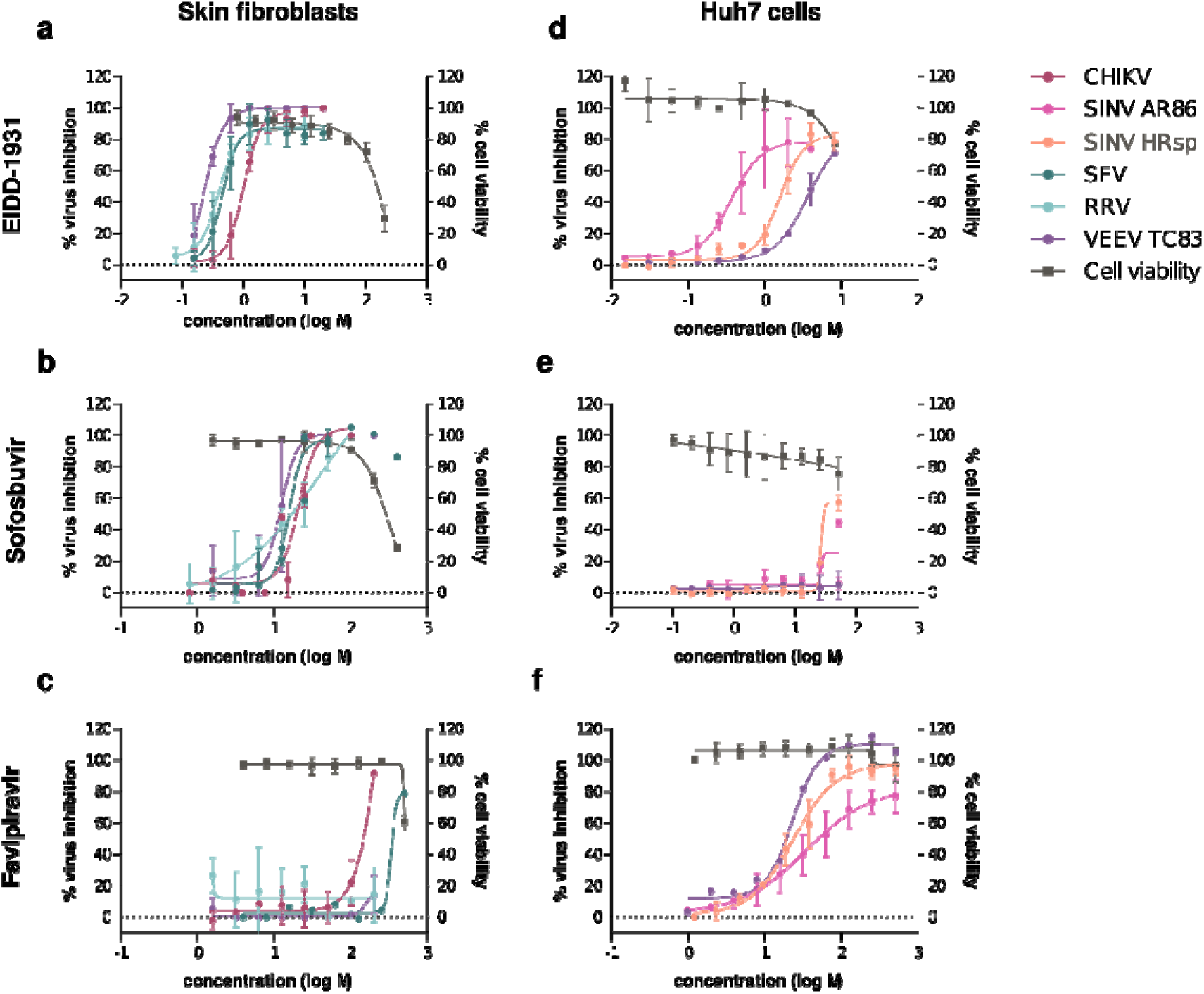
Antiviral efficacy of single compounds against alphaviruses in human cells. Dose-response activity of (a,d) EIDD-1931 (MPV), (b,e) SOF and (c,f) FAV against cytopathic effect induced by CHIKV, SFV, VEEV, RRV or SINV (AR86 and HRsp strain), quantified in (a-c) skin fibroblasts or (d-f) Huh7 cells by the MTS or ATP method at 72 hours post infection. Data represent percentage of virus inhibition (left y-axis), or cell viability (right y-axis) compared to virus or cell control samples, respectively, and is shown as mean values ± standard deviation from at least three independent experiments.

In Huh7 cells, MPV again showed potent pan-alphavirus activity with EC_50_ values in the sub to low single-digit micromolar range (Fig 1d, Table 2). Treatment with SOF did not inhibit VEEV TC83 and SINV AR86 in this cell line (Fig 1e). FAV was active against SINV HRsp, SINV AR86 and VEEV TC83 (Fig 1f, Table 2). Collectively, these data indicate that three approved RNA polymerase inhibitors block the replication of 6 different alphaviruses; however, the antiviral activity of FAV and SOF was virus- and cell type dependent.

### Identification of synergistic drug pairs against alphaviruses in human skin fibroblasts

Checkerboard combination assays (see Methods) were then performed with combinations of MPV (active metabolite EIDD-1931), SOF and FAV against CHIKV in human skin fibroblast cells (Fig 2). The degree of synergistic inhibition was defined as potency above that predicted by the multiplicative Bliss independence model. Previous studies have indicated that synergy scores >10 may be indicative of biologically meaningful combinatorial effects.^19^ Combination treatment with SOF and FAV resulted in an overall mean Bliss score of 14·3, demonstrating significant synergistic suppression of CHIKV (Fig 2a, C). A pronounced score of 33·1 was observed in the most synergistic area (MSA; a 3 x 3 submatrix of drug concentrations where maximum synergy is observed) (Fig 2b). Interestingly, this MSA was observed in the low concentration ranges of the drugs (Fig 2c). The combination of SOF and MPV was associated with a mean Bliss score of 10·5, suggesting moderate synergistic antiviral activity. However, marked synergism occurred at low concentration ranges, with a mean MSA of 25·7 (Fig 2d). In contrast, the combination of FAV and MPV conferred a mean Bliss score of 3·7 suggesting that there was minimal synergistic antiviral activity (Fig 2e). However, synergism occurred at certain drug concentrations with a MSA of 15·5. No to minimal cytotoxicity was observed after combination treatment of uninfected cells across all combined concentrations, as assessed using SynToxProfiler (Fig S2).

**Figure 2.**
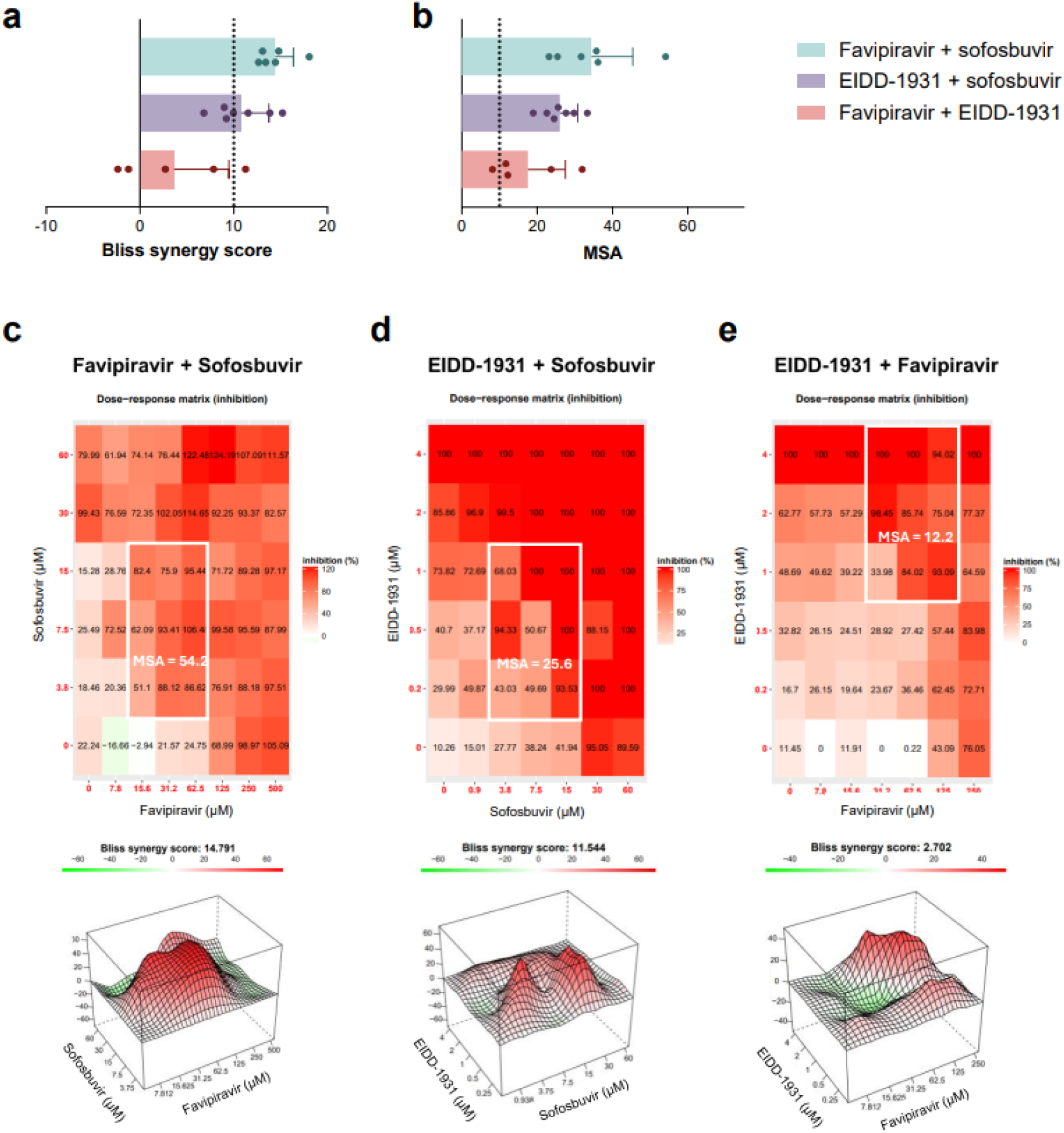
Antiviral efficacy of combination treatment against CHIKV in human skin fibroblasts. Antiviral efficacy of combinations of EIDD-1931 (MPV) and FAV, EIDD-1931 (MPV) and SOF or FAV and SOF against CHIKV in human skin fibroblasts. Inhibition of virus-induced CPE was quantified by the MTS method at 72 hours post infection. Data were analysed and visualized using

Next, it was investigated whether the synergistic activity of DAA combinations extended to other alphaviruses. FAV and SOF conferred similar synergistic suppression of SFV in skin fibroblasts, with a Bliss score of 14·2 and MSA score of 36·8 (Fig 3). In contrast, the combination of MPV and SOF did not result in an overall synergistic antiviral effect (Bliss score of 2·3), but synergism occurred at certain combinations of concentrations (MSA score of 16·4). Similarly, FAV and MPV did not act in an overall synergistic manner against SFV (Bliss score of 7·5) but synergized at certain higher concentration ranges (MSA score of 22·6).

**Figure 3.**
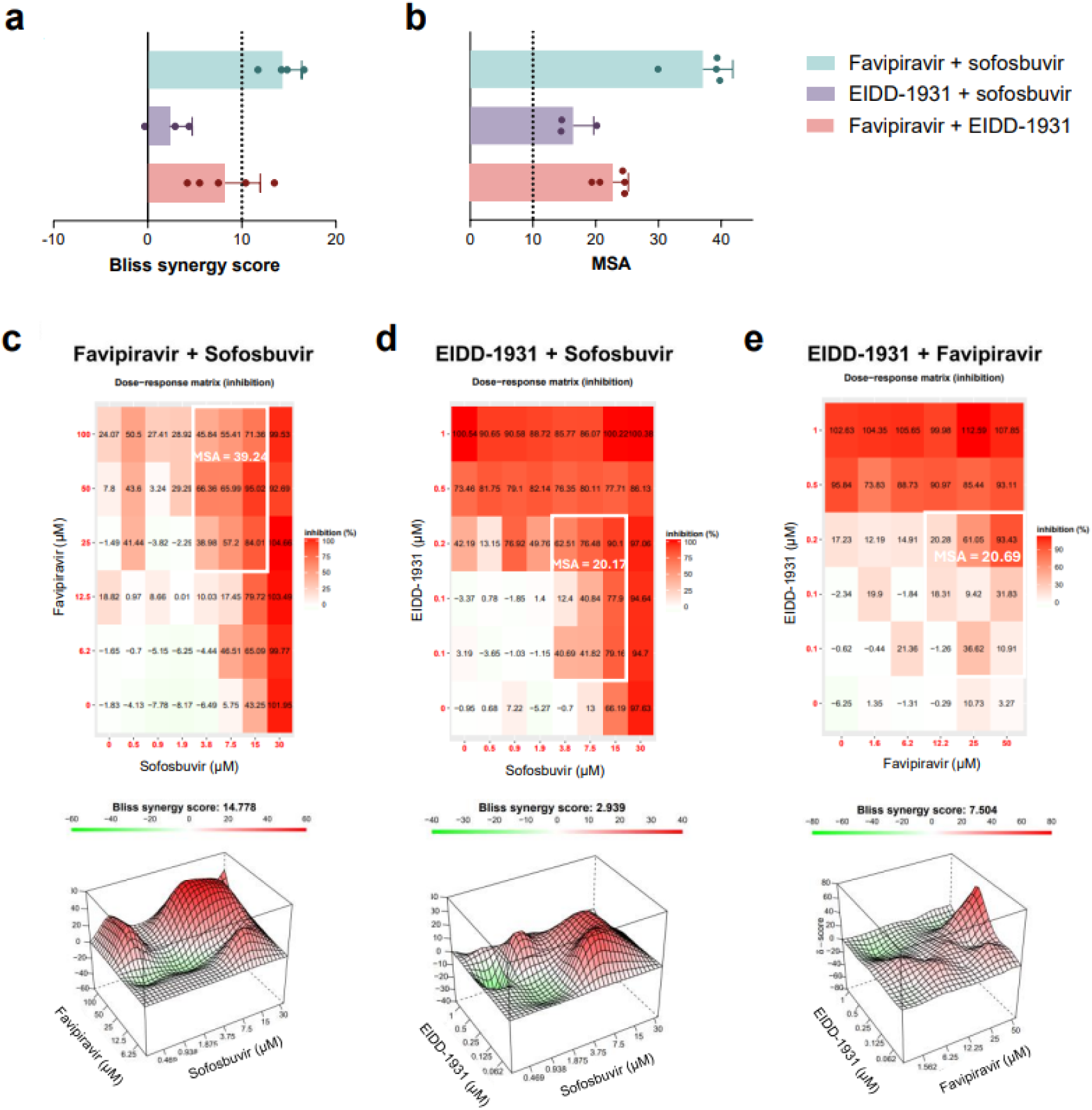
Antiviral efficacy of combination treatment against SFV in human skin fibroblasts. Antiviral efficacy of combinations of EIDD-1931 (MPV) and FAV, EIDD-1931 (MPV) and SOF or FAV and SOF against SFV in human skin fibroblasts. Inhibition of virus-induced CPE was quantified by the MTS method at 72 hours post infection. Data were analysed and visualized using

### DAA combinations confer additive to synergistic antiviral effects in Huh7 cells

To study whether these combinations also show synergistic activity in another human cell line that reflects the systemic infection associated with alphavirus infections, we next tested the DAA combinations against SINV and VEEV infection in Huh7 cells (Fig 4). MPV and FAV showed modest synergistic antiviral effects against VEEV TC83 with a mean Bliss score of 10·3 and mean MSA of 22·5 (Fig 4a). The MPV+FAV combination conferred additive effects against both SINV HRsp and SINV AR86, with mean Bliss scores of 9·0 and 9·4 and mean MSA scores of 14·4 and 18·0, respectively (Fig 4b, c). While SOF had modest antiviral efficacy as a solo agent against SINV HRsp and SINV AR86 in Huh7 cells (Fig 1), it synergized with FAV against these two viruses (Fig 4b, c). SOF did not significantly synergize with MPV against VEEV (Bliss score 1·8) and SINV HRsp (Bliss score 6·7) or AR86 (Bliss score 7·9) (Fig 4). An overview of overall Bliss synergy scores and MSAs for each specific virus and cell line is presented in Table S1. Plots highlighting the potency of each combination, as well as areas of synergy and antagonism across the full dose response matrix are depicted in Fig S3 and S4. Only minimal cytotoxicity was observed after combination treatment of uninfected cells, as analysed using SynToxProfiler (Fig S5).

**Figure 4.**
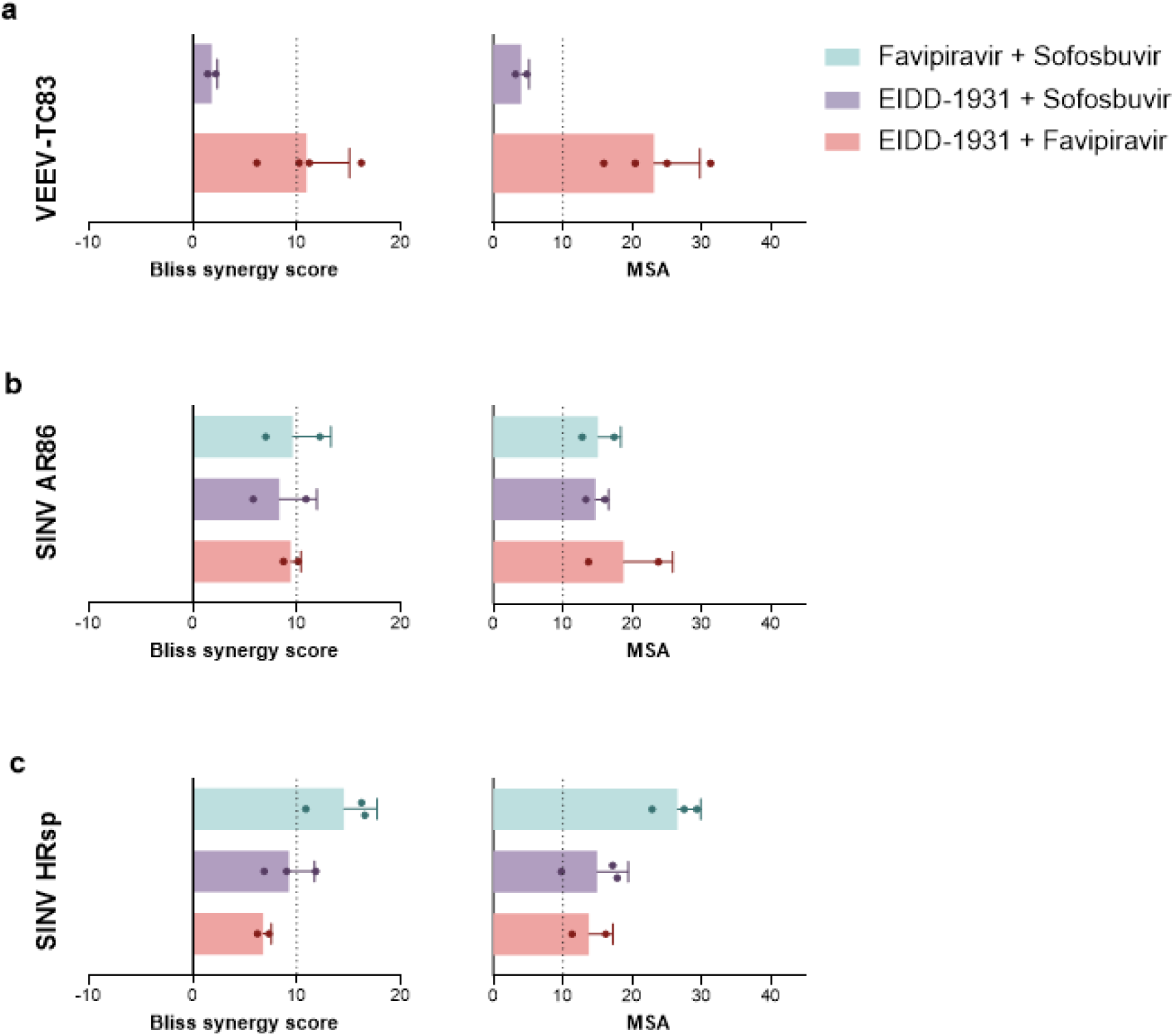
Antiviral efficacy of combination treatment against SINV and VEEV in Huh7 cells. Antiviral efficacy of combinations of EIDD-1931 (MPV) and FAV, EIDD-1931 (MPV) and SOF or FAV and SOF against (a) VEEV TC83, (b) SINV AR86, (c) SINV HRsp in Huh7 cells. Inhibition of virus-induced CPE was quantified by the ATP method at 72 hours post infection. Data were analysed and visualized using SynergyFinder 3·0 based on the Bliss independence model. Left panels: overall Bliss Synergy score averaged over all dose combination measurements of the matrix, representing the percentage excess response compared to the expected responses. Bliss Synergy scores lower than - 10, between -10 and 10 or larger than 10 indicate combinations which are likely to be antagonistic, additive or synergistic, respectively. Right panels: maximum Bliss synergy score representing the most synergistic 3-by-3 window in the full dose-response matrix (MSA). Data in panels a-c indicate the mean scores and standard deviations of 2-4 independent experiments (with each experiment denoted by a dot). MSA; most synergistic area.

### Combination DAA treatment enhanced the antiviral activity against CHIKV infection in mice

To explore the therapeutic potential and combined efficacy of DAAs *in vivo*, we next determined whether combining these drugs would increase the inhibitory effect against CHIKV in comparison to the respective monotherapies using a CHIKV infection model in AG129 mice. As EIDD-1931, the active metabolite of MPV, was identified as the most potent and broad-spectrum inhibitor in cell lines, we selected MPV for *in vivo* combination studies. To complement its antiviral activity, which is induced via lethal mutagenesis of the viral RNA genome, we combined MPV with SOF, which acts through chain termination, aiming to achieve synergistic antiviral effect through distinct mechanisms. To evaluate the efficacy of the combination, we sought to use suboptimal doses for each drug that resulted in ≤1·5 log10 reduction in the joint infectious virus titres (TCID50/mg joint tissue). We first evaluated the dose-response efficacy of each drug as a monotherapy in CHIKV-infected AG129 mice (Fig 5). For MPV, oral administration of 100, 50 and 10 mg/kg doses was initiated prior to subcutaneous infection with CHIKV (100 PFU) and administered for three consecutive days. On day 2 pi, treatment with 100 and 50 mg/kg of MPV significantly reduced foot swelling by 20% and 22% compared to vehicle-treated mice (Fig 5a). By day 3 pi, swelling was almost entirely prevented in the higher-dose groups, with reductions of 62% (100 mg/kg) and 55% (50 mg/kg), whereas the low dose of 10 mg/kg did not significantly improve swelling. At day 2 pi, all doses of MPV suppressed infectious CHIKV in mouse serum while at day 3 pi, MPV conferred dose-dependent decreases in infectious virus in serum (Fig 5b). Viral RNA was also detectable in serum on day 3 pi and was significantly reduced by MPV in a dose-dependent manner (Fig 5e). In the left ankle, the site of initial infection, significant decreases in infectious virus and viral RNA were observed at the highest dose of MPV (Fig 5c, f). In the contralateral ankle, MPV induced dose-dependent decreases in infectious virus and viral RNA (Fig 5d, g). Comparable results were found in distant tissues, including the liver, brain, spleen and lymph node (Fig S6). Importantly, MPV treatment was well-tolerated by all mice, with no adverse effects across all doses (Fig S8).

**Figure 5.**
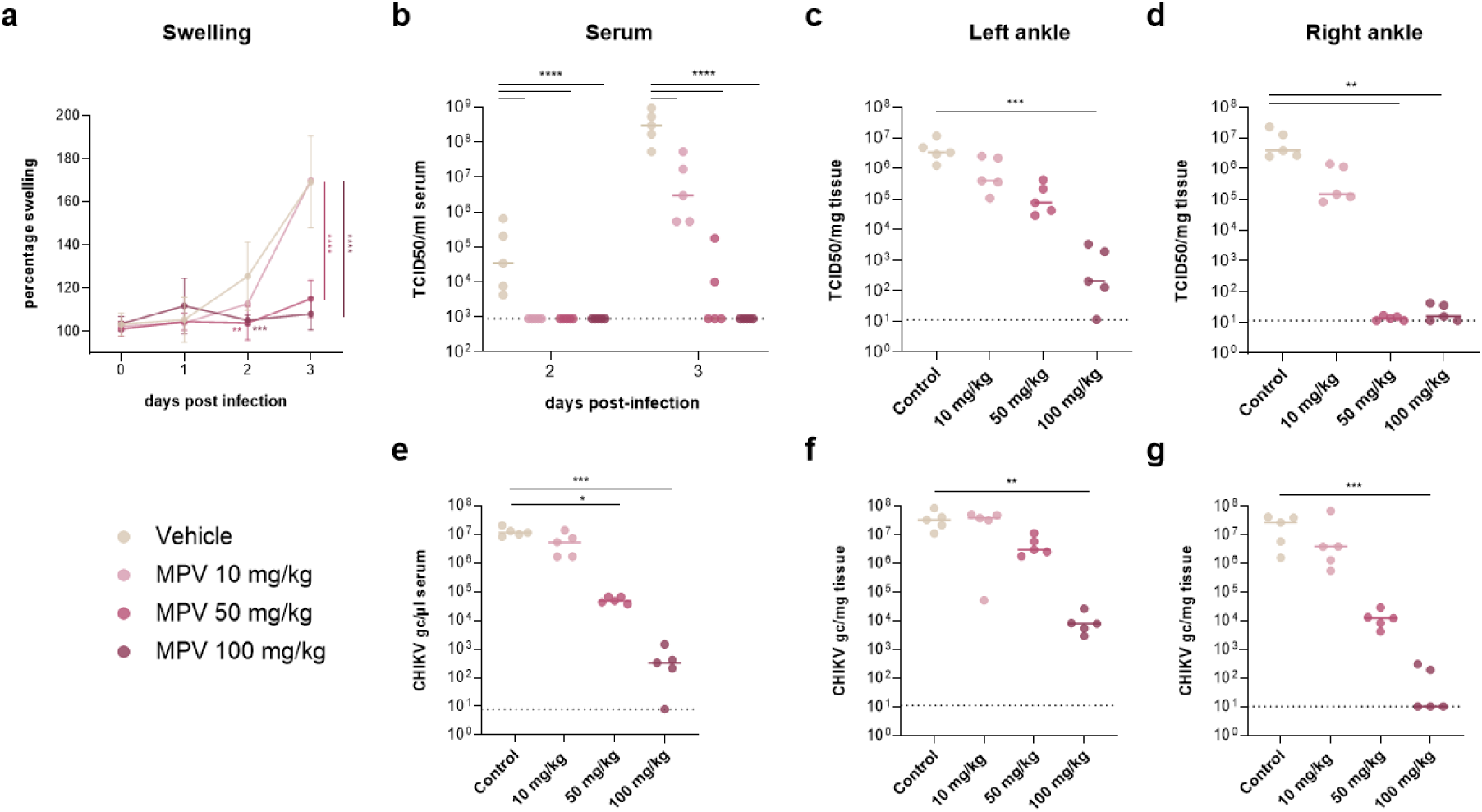
Dose-dependent efficacy of MPV against CHIKV infections AG129 mice. AG129 mice (n=5 per group) were treated orally with different single doses of MPV (10, 50, 100 mg/kg) and infected subcutaneously with CHIKV (100 PFU) in the left hind foot. (a) Median and standard deviation of percentage swelling of the infected foot relative to the contralateral foot, as measured daily using a digital calliper. (b-d) Infectious virus titres in tissues, quantified by end-point titrations on Vero cells. (e-g) Viral RNA levels in tissues, quantified by qRT-PCR. (b,e) Virus loads in blood collected by submandibular (day 2 pi) or cardiac (day 3 pi) puncture. (c,f) Virus loads in the left, infected ankle joint on day 3 pi. (d,g) Virus loads in the right, contralateral ankle joint on day 3 pi. Individual data points are shown, with solid lines representing median values. Statistical significance was assessed with (a,b) two-way repeated measures ANOVA with Tukey’s correction or (c-g) Kruskal-Wallis test with Dunn’s multiple comparisons test (*, p<0·05; **, p<0·01; ***, p<0·005; ****, p<0·0001). (b-d) Dotted lines represent the LOQ; (e-g) dotted lines represent the LOD. Gc, genome copies; TCID50, tissue culture infectious dose 50; pi, post infection; LOQ, limit of quantification; LOD, limit of detection.

We also evaluated the efficacy of SOF as a monotherapy against CHIKV in AG129 mice. Treatment with 40 and 80 mg/kg of SOF did not significantly improve CHIKV-induced foot swelling over the course of the infection (Fig 6a). Consistently, neither treatment group showed significant reductions in virus loads compared to the vehicle-treated group amongst all tissues examined (serum, lymph node, ankle joints, liver, spleen and brain) (Fig 6b-g, Fig S7).

**Figure 6.**
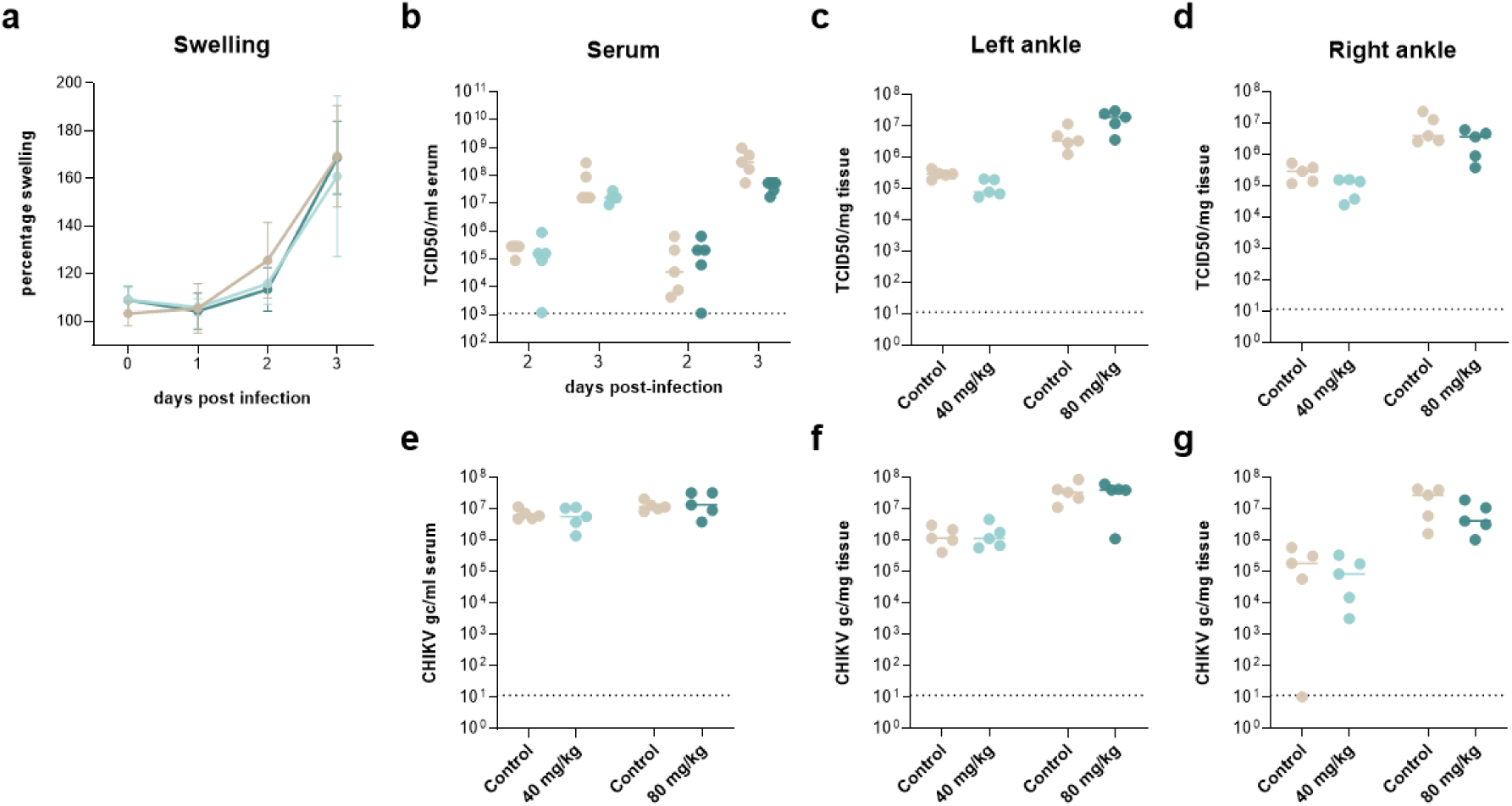
Dose-dependent efficacy of SOF against CHIKV infections in AG129 mice. AG129 mice (n=5 per group) were treated orally with different single doses of SOF (40, 80 mg/kg) and infected subcutaneously with CHIKV (100 PFU) in the left hind foot. (a) Median and standard deviation of percentage swelling of the infected foot relative to the contralateral foot, as measured daily using a digital calliper. (b-d) Infectious virus titres in tissues, quantified by end-point titrations on Vero cells. (e-g) Viral RNA levels in tissues, quantified by qRT-PCR. (b,e) Virus loads in blood collected by submandibular (day 2 pi) or cardiac (day 3 pi) puncture. (c,f) Virus loads in the left, infected ankle joint on day 3 pi. (d,g) Virus loads in the right, contralateral ankle joint on day 3 pi. Individual data points are shown, with solid lines representing median values. Statistical significance was assessed with (a,b) two-way repeated measures ANOVA with Tukey’s correction or (c-g) Kruskal-Wallis test with Dunn’s multiple comparisons test (ns, p>0·05). (b-d) Dotted lines represent the LOQ; (e-g) dotted lines represent the LOD. Gc, genome copies; TCID50, tissue culture infectious dose 50; pi, post infection; LOQ, limit of quantification; LOD, limit of detection.

Informed by these monotherapy experiments, we next evaluated the combined efficacy of MPV with SOF. Mice were treated either with suboptimal doses of 80 mg/kg of SOF, 10 mg/kg of MPV or a combination of SOF and MPV (80 + 10 mg/kg) (Fig 7). Monotherapy with MPV or SOF significantly reduced foot swelling on day 2 pi by 19% or 25%, respectively (Fig 7a). The combination therapy resulted in a more significant reduction of 34% compared to the vehicle treatment. By day 3 pi, SOF alone showed no reduction of swelling, while swelling decreased by 7·8% upon monotherapy with MPV. Interestingly, a marked reduction of 18% was achieved in mice treated with the combination, exceeding the expected additive effect of the individual treatments. A similar trend was observed for infectious virus titres in serum, which were reduced by 0·2 log_10_ and 1 log_10_ TCID_50_/mL upon single treatment with SOF and MPV, respectively, while the combination therapy resulted in a substantial decrease of 2 log_10_ TCID_50_/ml on day 3 pi (Fig 7b), which was significantly different from both monotherapies. In both ankle joints, infectious virus titres were reduced more significantly following combination treatment compared to treatment with single compounds, with reductions of 1·4 log_10_ TCID_50_/mg in the ipsilateral ankle and 2·0 log_10_ TCID_50_/mg in the contralateral ankle (Fig 7c,d). Consistently, viral RNA loads were reduced by 1·3 log_10_ genome copies (gc) following combination therapy, compared to only modest changes in the monotherapy groups (MPV: decrease of 0·2 log_10_, SOF increase of 0·1 log_10_ gc/mg tissue) (Fig 7e). Similarly, in the contralateral ankle, combination treatment resulted in a 0·9 log_10_ reduction in viral RNA, outperforming the individual treatments (SOF: increase of 0·2 log_10_ gc, MPV: decrease of 0·5 log_10_ gc) (Fig 7f). However, the combination did not outperform the efficacy of the medium (50 mg/kg) and high (100 mg/kg) dose of MPV monotherapy (Fig 5). Crucially, no weight loss or adverse effects were observed in any of the treatment groups (Fig S8). These data demonstrate the potential for enhanced antiviral efficacy of the combined use of MPV and SOF against CHIKV *in vivo*.

**Figure 7.**
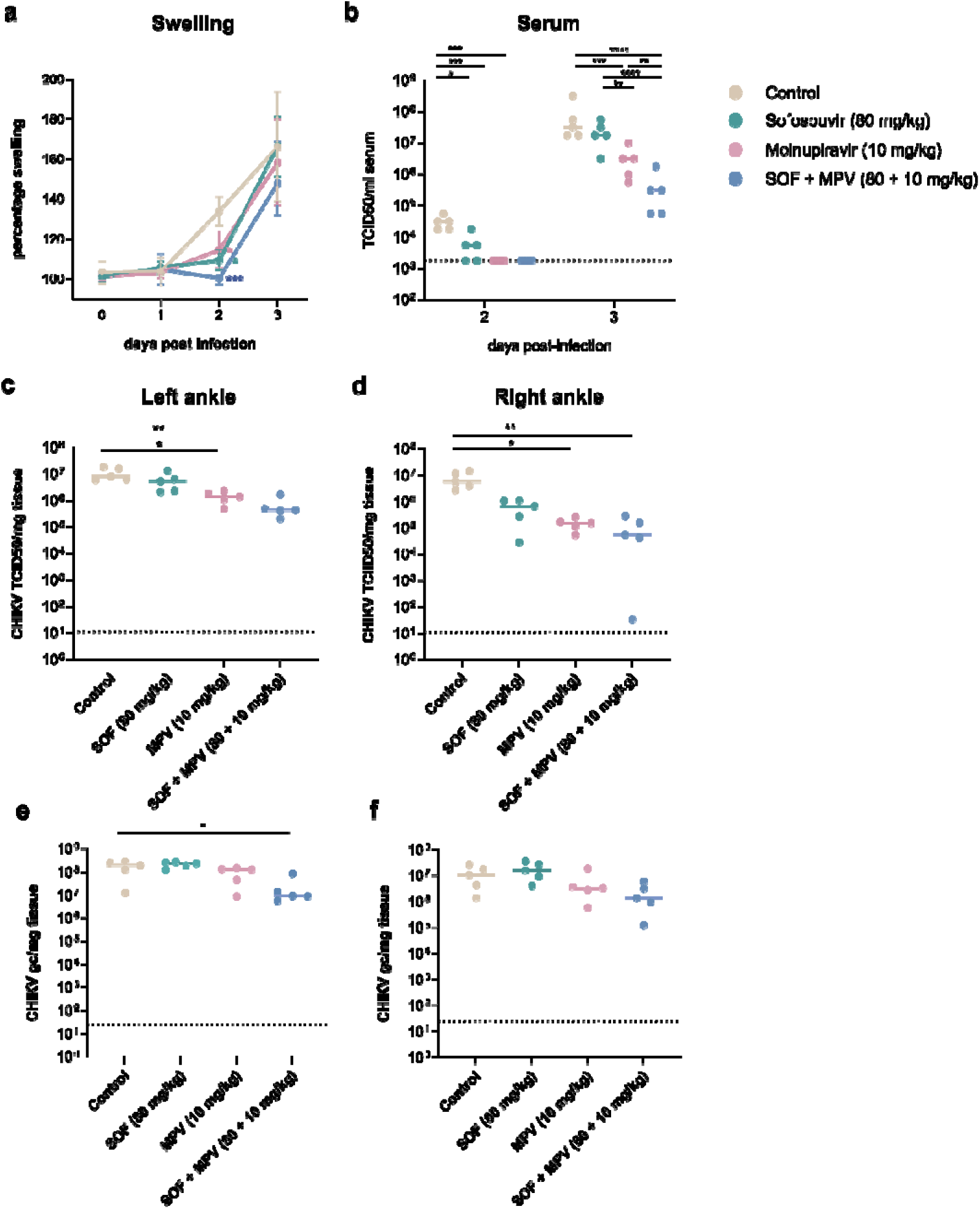
Combined efficacy of MPV with SOF against CHIKV infections in AG129 mice. AG129 mice (n=5 per group) were treated orally with either single doses of MPV (10 mg/kg), SOF (80 mg/kg) or a combination of MPV and SOF (10 + 80 mg/kg) and infected subcutaneously with CHIKV (100 PFU) in the left hind foot. (a) Median and standard deviation of percentage swelling of the infected foot relative to the contralateral foot, as measured daily by using a digital calliper. (b-d) Infectious virus titres in tissues, quantified by end-point titrations on Vero cells. (e-f) Viral RNA levels in tissues, quantified by qRT-PCR. (b) Virus loads in blood collected by submandibular (day 2 pi) or cardiac (day 3 pi) puncture. (c,e) Virus loads in the left, infected ankle joint on day 3 pi. (d,f) Virus loads in the right, contralateral ankle joint on day 3 pi. Individual data points are shown, with solid lines representing median values. Statistical significance was assessed with (a,b) two-way repeated measures ANOVA with Tukey’s correction or (c-f) Kruskal-Wallis test with Dunn’s multiple comparisons test (*, p<0·05; **, p<0·01; ***, p<0·005). (b-d) Dotted lines represent the LOQ; (e-f) dotted lines represent the LOD. Gc, genome copies; TCID50, tissue culture infectious dose 50; pi, post infection; LOQ, limit of quantification; LOD, limit of detection.

### Viral genome mutational profiles upon MPV and SOF treatment of CHIKV-infected mice

MPV is known to act as a mutagenic drug against other RNA viruses.^65^ To determine whether this mechanism of action also applies to CHIKV, we performed sequencing on viral RNA isolated from the joints of CHIKV-infected mice to profile mutations upon treatment with MPV (10, 50 and 100 mg/kg) and SOF (80 mg/kg). Treatment with MPV resulted in a dose-dependent increase in the overall mutation count in the CHIKV genome (Fig 8b), with the largest increase in T-to-C transitions (Fig 8a, c). One sample of the MPV 10 mg/kg group and two samples of the MPV 100 mg/kg group had to be excluded due to low read counts. In contrast to MPV, SOF showed minimal mutagenic activity, with a mutation profile comparable to vehicle-treated mice (Fig 8b). One sample in the SOF group showed unexpectedly high mutation counts; this outlier was excluded for further analysis (Fig 8a).

**Figure 8.**
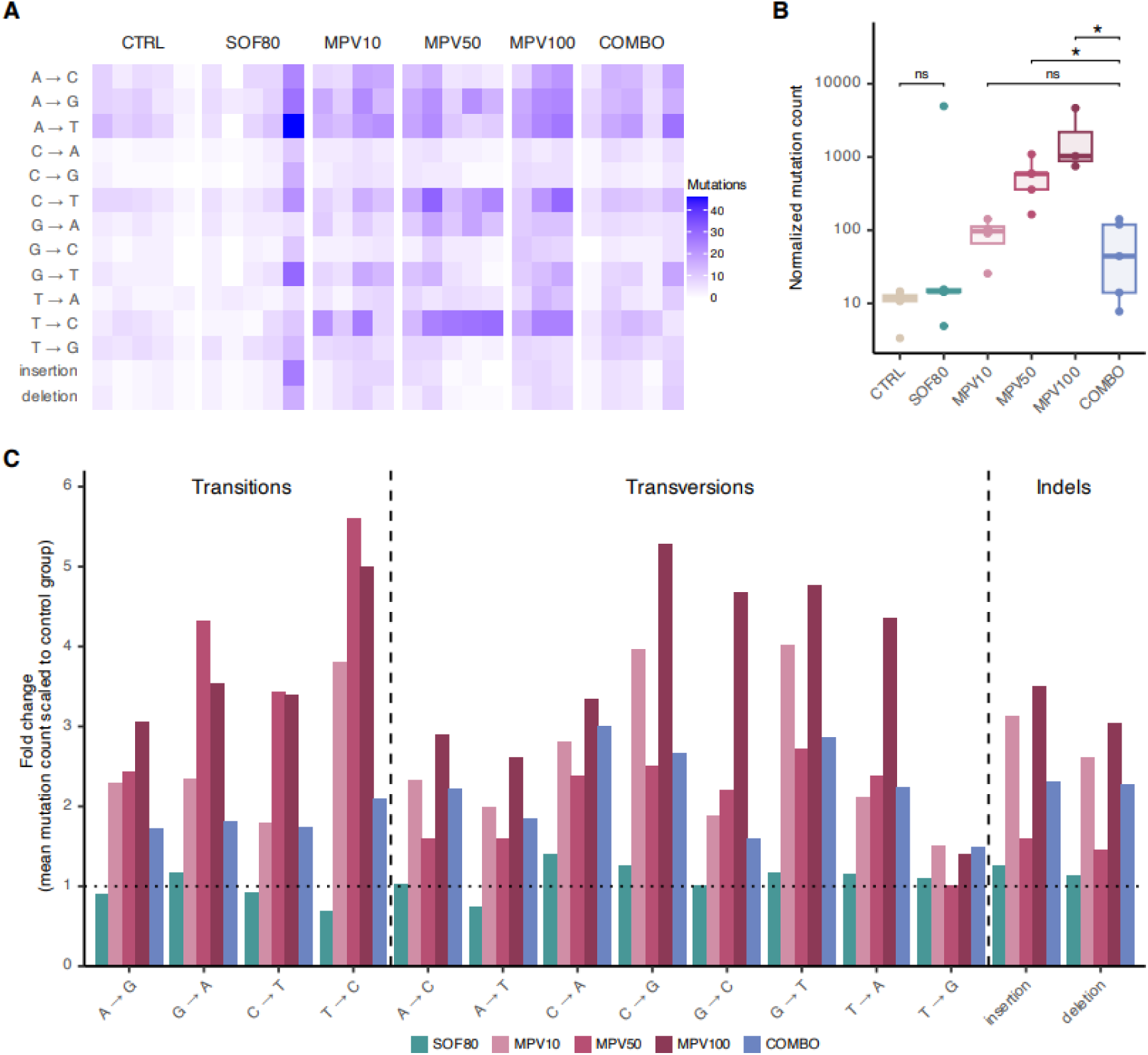
CHIKV mutation profiles of MPV- and SOF-treated mice. Sequencing was performed on viral RNA isolated from contralateral ankle joints of CHIKV-infected AG129 mice (n=5 per condition) that were treated with vehicle, MPV (10, 50, 100 mg/kg), SOF (80 mg/kg) or a combination of MPV (10 mg/kg) and SOF (80 mg/kg). (a) Mutation profile of viral RNA across treatment groups. The colour intensity represents the mutation count per genome for each specific mutation type. (b) Total mutation count in viral RNA normalized to the total number of reads per sample. Data is represented as boxplots, showing individual data points with solid lines indicating the median values. Statistical significance was assessed with a one-sided Wilcoxon test with Benjamini-Hochberg correction. *, p<0·05. (c) Bar plot showing the fold change in the mean count of each mutation type for each treatment group (n=5) relative to the control group (vehicle treatment). The dotted line indicates no change in mutation count compared to the vehicle-treated control group.

The total mutation count was significantly lower upon combination therapy compared to single treatment with MPV doses of 50 and 100 mg/kg (Fig 8b). In particular, T-to-C, C-to-T, and G-to-A transition mutations were significantly lower upon combination therapy (Table S2). Moreover, 2 out of 5 samples from mice treated with the MPV + SOF combination exhibited lower mutation counts than the MPV 10 group and were comparable to the vehicle- and SOF-treated group (Fig 8b).

## Discussion

Combination therapy using two or three DAAs has proven key in the successful treatment of chronic RNA virus infections such as HIV and HCV.^17,18^ Recently, this approach has also shown promise in combating pandemic viruses. For example, the combination of MPV with another nucleoside analogue, remdesivir, was shown to reduce the replication of SARS-CoV-2 in hamsters.^22^ Similarly, MPV enhanced the potential of FAV to prevent replication and transmission of SARS-CoV-2 compared to monotherapy.^66^ Because of the current absence of antiviral drugs to treat alphavirus-induced diseases and the significant morbidity linked to alphavirus infections, it is important to evaluate the efficacy of drug combinations. The use of existing approved drugs circumvents many of the early hurdles of drug discovery and also provides a promising avenue for rapidly addressing the urgent need for anti-alphavirus therapeutics. The viral RNA-dependent RNA polymerase (RdRp), the target of nucleoside analogue drugs, is encoded by the alphavirus nsP4 protein. Given this is a highly conserved protein amongst alphaviruses and other ssRNA viruses, it presents great potential for broad-spectrum activity. SOF has been shown to inhibit the RdRp of several RNA viruses and modeling studies have predicted that SOF engages CHIKV’s nsP4.^61^ Moreover, it was shown that SOF could inhibit CHIKV infections in human astrocytes and liver cells.^61^ The incorporation of SOF into the viral RNA genome by its viral RdRps results in chain termination and consequently prevents further viral replication.^67^ Here, we showed that SOF has broad anti-alphavirus activity in human skin fibroblasts. However, SOF showed minimal to no activity against VEEV and SINV in Huh7 cells. Additionally, we did not observe significant improvements upon SOF treatment of CHIKV-infected AG129 mice at doses of 80 mg/kg over a course of 4 days. A previously published mouse study reported that 40-80 mg/kg/day doses improved survival of CHIKV-infected neonatal mice and a dose of 20 mg/kg/day slightly reduced foot swelling and viral RNA in the joint (<1 log gc/ml) of adult Swiss mice.^61^ These different results can possibly be attributed to the different mouse models or drug administration routes that were used, the different time points of investigations, or the small sample sizes.

Additional studies will be required to determine why SOF fails to efficiently suppress VEEV and SINV in Huh-7 cells. It is possible that intracellular conversion of SOF to the triphosphate form (required for incorporation of SOF into the nascent RNA and chain termination of RNA replication) is reduced in Huh7 cells. However, this seems unlikely since Huh7 cells and various derivatives of this cell line have been widely used to study the antiviral efficacy of SOF versus multiple viruses including HCV,^68^ hepatitis A virus,^65^ and Zika virus.^69^ Despite its reduced overall activity in Huh7 cells, combining SOF with another DAA increased its antiviral activity compared to SOF as a monotherapy in Huh7 (Fig 4).

The combination of FAV and SOF showed consistent synergistic effects against all alphaviruses tested in two different human cell lines. Furthermore, the MSAs were seen in the low concentration ranges, suggesting that this combination could be more potent *in vivo* than FAV + MPV. FAV is recognized by the viral RdRp as a pseudopurine and can be incorporated in the viral RNA for an adenosine and a guanosine, whereas SOF is a uridine analogue. Therefore, they should not compete with each other to be incorporated. Moreover, both drugs act differently upon incorporation into the viral genome (inducing lethal mutagenesis and chain termination, respectively), which might explain their synergistic effects in cell culture. In contrast, the combination of MPV and FAV conferred modest synergistic effects *in vitro* against SINV, VEEV, and SFV, but not against CHIKV. In addition, the MSAs for MPV and FAV were observed only in the high concentration ranges, indicating that synergy was only achieved at high concentrations of both drugs. This suggests that MPV and FAV may be a less optimal drug combination. This might be because these drugs have similar mechanisms of action (i.e., error catastrophe). For the combination of MPV and SOF, modest combination effects were observed against CHIKV and SINV, but not against SFV and VEEV. For VEEV, the absence of synergism for MPV and SOF could be explained by the lack of antiviral activity of SOF in Huh7 cells against this virus. Additional drug combinations should be evaluated for inhibition of VEEV and related encephalitic alphaviruses. For CHIKV and SINV, the MSAs for MPV and SOF were observed in the low concentration ranges, suggesting that combining lower doses of both drugs may be sufficient to achieve antiviral efficacy. In addition, this drug combination demonstrated enhanced antiviral efficacy in CHIKV-infected mice, when compared to the respective monotherapies of MPV and SOF (Fig 7). However, higher doses of MPV as a monotherapy were still effective at reducing CHIKV viral loads and foot swelling in infected mice. A potential downside with MPV as a solo drug is its mutagenic action which creates the potential for generating antiviral drug-resistant, transmissible viruses.^70^ Thus, the combination regimen of MPV and SOF still requires optimization.

Lethal mutagenesis is a known mechanism of the antiviral activity of MPV against other RNA viruses.^28,71^ Because of this, concerns have been raised that the increased mutation rate upon MPV treatment could lead to the emergence of virus variants that are drug-resistant or have an increased fitness.^72^ We therefore performed deep sequencing of CHIKV genomes from contralateral joint tissues, revealing a substantial increase in mutation frequency following MPV treatment. The dose-dependent increase in the overall mutation count can explain the potent and dose-dependent reduction of virus loads that we observed in MPV-treated mice. It also confirms that MPV acts as a potent mutagen for CHIKV, where the accumulation of mutations will be deleterious for the virus and lead to non-viable progeny. Importantly, in combination with SOF, the mutation rate of CHIKV was reduced compared to single MPV treatment with the medium and high dose tested. This suggests that a suboptimal dose of MPV could be used in a combination to reduce virus infection and, at the same time, limit mutagenesis of remaining viral RNA. It was shown previously that the combination of nirmatrelvir and MPV resulted in improved efficacy against SARS-CoV-2, while nirmatrelvir was able to reduce the mutagenicity induced by MPV.^73^ This shows that optimized drug combinations containing a mutagenic drug could, in addition to enhancing antiviral activity, result in a substantial reduction of the mutation frequency, reducing the likelihood for emergence of drug-resistant variants.

While DAAs are the first choice to evaluate as combinations (as done here), testing combinations of DAAs with drugs that target host functions (Host Targeting Antivirals, HTAs) is also worthwhile. We and others showed that DAAs and HTAs are synergistic against SARS-CoV-2 in lung cells,^24,27^ and one DAA-HTA pair tested in a SARS-CoV-2 mouse model showed clear benefits.^27^ For alphaviruses, little is known about DAA-HTA combinations in relevant cells and mouse models and so future studies should consider this.

In conclusion, our study demonstrates the potential of combining approved oral nucleoside analogues as a strategy to combat alphavirus infections. By combining two drugs, we demonstrated enhanced antiviral activity in relevant human cell lines, as well as reduced virus dissemination and improved disease symptoms in an *in vivo* CHIKV mouse infection model. Furthermore, the combination regimen led to a reduced number of mutations that were induced in the viral genome, compared to single treatment with several higher doses of MPV. With these findings, we highlight the importance of combination therapies in overcoming the limitations of monotherapies and underscore the value of using existing antiviral drugs as a rapid and cost-effective strategy to address the unmet medical need for anti-alphavirus therapies. Importantly, drug combinations can serve as a critical tool for better preparedness in future outbreaks, where early therapeutic intervention will be essential to reduce disease burden and interrupt transmission cycles. Moving forward, optimizing these combinations for clinical application and exploring additional synergistic therapies will be key to enhancing global readiness against the ever-growing threat of alphaviruses.

## Contributors

Conceptualisation: SV, LD, SJP, JLH, JTS, JMW; Data curation: SV, LD, SJP; Formal analysis: SV, SJP, LDC, JW, EGA, RA; Funding acquisition: SJP, LD, JLH, JTS, JMW, JN; Investigation: SV, JW, EGA, LDC, RA; Methodology: SV, JW, EGA, LDC, RA, JM, SJP, LD; Project administration: LD, SJP; Resources: LD, SJP, JLH, JM, JN; Supervision: LD, SJP; Validation: SV, LD, SJP; Visualisation: SV, LD, SJP, LDC; Writing – original draft: SV, LD, SJP, LDC; Writing – review & editing: SV, LD, SJP, RA, JLH, JTS, JMW, JM, JN.

## Acknowledgements

This work was supported by a PhD fellowship granted to SV by the Research Foundation – Flanders (FWO) (11D5923N). LDC was also supported by Research Foundation – Flanders (FWO) PhD fellowship (11L1325N). SJP and JTS are partially supported by R01AI121129. We would like to thank the Rega Animalium staff for their support in the animal facility; Carolien De Keyzer and Thibault Francken for their help in animal experiments; Kristien Minner for assistance in antiviral assays; Xin Zhang for advice on drug combination studies. We also thank Karen Youngblood, Ariana Paul and Gillian Milstein for technical assistance. We thank Drs. Ilya Frolov and Mark Heise for virus reagent.

## References

1 Bartholomeeusen K, Daniel M, LaBeaud DA, Gasque P, Peeling RW, Stephenson KE, et al. Chikungunya fever. Nat Rev Dis Primers 2023; 9: 17.

2 de Lima Cavalcanti TYV, Pereira MR, de Paula SO, Franca RF de O. A Review on Chikungunya Virus Epidemiology, Pathogenesis and Current Vaccine Development. Viruses 2022; 14. doi:10.3390/V14050969.

3 Levi LI, Vignuzzi M. Arthritogenic Alphaviruses: A Worldwide Emerging Threat? Microorganisms 2019; 7: 133.

4 Vidal ERN, Frutuoso LCV, Duarte EC, Peixoto HM. Epidemiological burden of Chikungunya fever in Brazil, 2016 and 2017. Tropical Medicine & International Health 2022; 27: 174–184.

5 Rodríguez-Morales AJ, Simon F. Chronic chikungunya, still to be fully understood. International Journal of Infectious Diseases 2019; 86: 133–134.

6 Montalvo Zurbia-Flores G, Reyes-Sandoval A, Kim YC. Chikungunya Virus: Priority Pathogen or Passing Trend? Vaccines (Basel) 2023; 11. doi:10.3390/VACCINES11030568.

7 Guerrero-Arguero I, Tellez-Freitas CM, Weber KS, Berges BK, Robison RA, Pickett BE. Alphaviruses: Host pathogenesis, immune response, and vaccine & treatment updates. J Gen Virol 2021; 102. doi:10.1099/JGV.0.001644.

8 Kurkela S, Manni T, Vaheri A, Vapalahti O. Causative Agent of Pogosta Disease Isolated from Blood and Skin Lesions. Emerg Infect Dis 2004; 10: 889.

9 Baxter VK, Heise MT. Immunopathogenesis of alphaviruses. Adv Virus Res 2020; 107: 315.

10 Contu L, Balistreri G, Domanski M, Uldry AC, Mühlemann O. Characterisation of the Semliki Forest Virus-host cell interactome reveals the viral capsid protein as an inhibitor of nonsense-mediated mRNA decay. PLoS Pathog 2021; 17: e1009603.

11 Weaver SC, Ferro C, Barrera R, Boshell J, Navarro JC. Venezuelan equine encephalitis. Annu Rev Entomol 2004; 49: 141–174.

12 Ronca SE, Dineley KT, Paessler S. Neurological sequelae resulting from encephalitic alphavirus infection. Front Microbiol 2016; 7: 198666.

13 Hawley RJ, Eitzen J. Biological weapons - A primer for microbiologists. Annu Rev Microbiol 2001; 55: 235–253.

14 Abdelnabi R, Delang L. Antiviral Strategies against Arthritogenic Alphaviruses. Microorganisms 2020; 8: 1–18.

15 August A, Attarwala HZ, Himansu S, Kalidindi S, Lu S, Pajon R et al. A phase 1 trial of lipid-encapsulated mRNA encoding a monoclonal antibody with neutralizing activity against Chikungunya virus. Nat Med 2021; 27: 2224.

16 Weber WC, Streblow DN, Coffey LL. Chikungunya Virus Vaccines: A Review of IXCHIQ and PXVX0317 from Pre-Clinical Evaluation to Licensure. BioDrugs 2024; 38. doi:10.1007/S40259-024-00677-Y.

17 Naggie S, Muir AJ. Oral Combination Therapies for Hepatitis C Virus Infection: Successes, Challenges, and Unmet Needs. Annu Rev Med 2017; 68: 345–358.

18 Gibas KM, Kelly SG, Arribas JR, Cahn P, Orkin C, Daar ES et al. Two-drug regimens for HIV treatment. Lancet HIV 2022; 9: e868.

19 Herring S, Oda JM, Wagoner J, Kirchmeier D, O’Connor A, Nelson EA et al. Inhibition of Arenaviruses by Combinations of Orally Available Approved Drugs. Antimicrob Agents Chemother 2021; 65. doi:10.1128/AAC.01146-20.

20 Dyall J, Nelson EA, Dewald LE, Guha R, Hart BJ, Zhou H et al. Identification of Combinations of Approved Drugs With Synergistic Activity Against Ebola Virus in Cell Cultures. J Infect Dis 2018; 218: S672–S678.

21 Finch CL, Dyall J, Xu S, Nelson EA, Postnikova E, Liang JY, et al. Formulation, Stability, Pharmacokinetic, and Modeling Studies for Tests of Synergistic Combinations of Orally Available Approved Drugs against Ebola Virus In Vivo. Microorganisms 2021; 9: 1–21.

22 Abdelnabi R, Maes P, de Jonghe S, Weynand B, Neyts J. Combination of the parent analogue of remdesivir (GS-441524) and molnupiravir results in a markedly potent antiviral effect in SARS-CoV-2 infected Syrian hamsters. Front Pharmacol 2022; 13. doi:10.3389/FPHAR.2022.1072202.

23 Do TND, Abdelnabi R, Boda B, Constant S, Neyts J, Jochmans D. The triple combination of Remdesivir (GS-441524), Molnupiravir and Ribavirin is highly efficient in inhibiting coronavirus replication in human nasal airway epithelial cell cultures and in a hamster infection model. Antiviral Res 2024; 231. doi:10.1016/J.ANTIVIRAL.2024.105994.

24 Wagoner J, Herring S, Hsiang T-Y, Ianevski A, Biering SB, Xu S et al. Combinations of Host- and Virus-Targeting Antiviral Drugs Confer Synergistic Suppression of SARS-CoV-2. Microbiol Spectr 2022; 10. doi:10.1128/SPECTRUM.03331-22.

25 White JM, Schiffer JT, Bender Ignacio RA, Xu S, Kainov D, Ianevski A et al. Drug Combinations as a First Line of Defense against Coronaviruses and Other Emerging Viruses. mBio 2021; 12. doi:10.1128/MBIO.03347-21.

26 Rosenke K, Lewis MC, Feldmann F, Bohrnsen E, Schwarz B, Okumura A et al. Combined molnupiravir-nirmatrelvir treatment improves the inhibitory effect on SARS-CoV-2 in macaques. JCI Insight 2023; 8. doi:10.1172/JCI.INSIGHT.166485.

27 Schultz DC, Johnson RM, Ayyanathan K, Miller J, Whig K, Kamalia B et al. Pyrimidine inhibitors synergize with nucleoside analogues to block SARS-CoV-2. Nature 2022; 604: 134–140.

28 Li P, Wang Y, Lavrijsen M, Lamers MM, de Vries AC, Rottier RJ et al. SARS-CoV-2 Omicron variant is highly sensitive to molnupiravir, nirmatrelvir, and the combination. Cell Res 2022; 32: 322–324.

29 White JM, Schiffer JT, Bender Ignacio RA, Xu S, Kainov D, Ianevski A et al. Drug Combinations as a First Line of Defense against Coronaviruses and Other Emerging Viruses. mBio 2021; 12. doi:10.1128/MBIO.03347-21/ASSET/437CDE00-3D5C-47C4-849D-32FA9ECF7064/ASSETS/IMAGES/MEDIUM/MBIO.03347-21-F001.GIF.

30 Gallegos KM, Drusano GL, Dlllargenio DZ, Brown AN. Chikungunya Virus: In Vitro Response to Combination Therapy With Ribavirin and Interferon Alfa 2a. J Infect Dis 2016; 214: 1192.

31 Rothan HA, Bahrani H, Mohamed Z, Teoh TC, Shankar EM, Rahman NA et al. A combination of doxycycline and ribavirin alleviated chikungunya infection. PLoS One 2015; 10. doi:10.1371/JOURNAL.PONE.0126360.

32 Rothan HA, Bahrani H, Abdulrahman AY, Mohamed Z, Teoh TC, Othman S et al. Mefenamic acid in combination with ribavirin shows significant effects in reducing chikungunya virus infection in vitro and in vivo. Antiviral Res 2016; 127: 50–56.

33 Briolant S, Garin D, Scaramozzino N, Jouan A, Crance JM. In vitro inhibition of Chikungunya and Semliki Forest viruses replication by antiviral compounds: synergistic effect of interferon-α and ribavirin combination. Antiviral Res 2004; 61: 111–117.

34 Lu JW, Hsieh PS, Lin CC, Hu MK, Huang SM, Wang YM et al. Synergistic effects of combination treatment using EGCG and suramin against the chikungunya virus. Biochem Biophys Res Commun 2017; 491: 595–602.

35 Franco EJ, Tao X, Hanrahan KC, Zhou J, Bulitta JB, Brown AN. Combination Regimens of Favipiravir Plus Interferon Alpha Inhibit Chikungunya Virus Replication in Clinically Relevant Human Cell Lines. Microorganisms 2021; 9: 1–16.

36 Furuta Y, Gowen BB, Takahashi K, Shiraki K, Smee DF, Barnard DL. Favipiravir (T-705), a novel viral RNA polymerase inhibitor. Antiviral Res 2013; 100: 446–454.

37 Maas BM, Strizki J, Miller RR, Kumar S, Brown M, Johnson MG et al. Molnupiravir: Mechanism of action, clinical, and translational science. Clin Transl Sci 2024; 17. doi:10.1111/CTS.13732.

38 Lawitz E, Jacobson IM, Nelson DR, Zeuzem S, Sulkowski MS, Esteban R et al. Development of sofosbuvir for the treatment of hepatitis C virus infection. Ann N Y Acad Sci 2015; 1358: 56– 67.

39 Molnupiravir | C13H19N3O7 | CID 145996610 - PubChem. https://pubchem.ncbi.nlm.nih.gov/compound/Molnupiravir (accessed 6 Feb2025).

40 Favipiravir | C5H4FN3O2 | CID 492405 - PubChem. https://pubchem.ncbi.nlm.nih.gov/compound/Favipiravir (accessed 6 Feb2025).

41 Sofosbuvir | C22H29FN3O9P | CID 45375808 - PubChem. https://pubchem.ncbi.nlm.nih.gov/compound/Sofosbuvir (accessed 6 Feb2025).

42 Couderc T, Chrétien F, Schilte C, Disson O, Brigitte M, Guivel-Benhassine F et al. A Mouse Model for Chikungunya: Young Age and Inefficient Type-I Interferon Signaling Are Risk Factors for Severe Disease. PLoS Pathog 2008; 4. doi:10.1371/JOURNAL.PPAT.0040029.

43 Ianevski A, Giri AK, Aittokallio T. SynergyFinder 3.0: an interactive analysis and consensus interpretation of multi-drug synergies across multiple samples. Nucleic Acids Res 2022; 50: W739–W743.

44 Ianevski A, Timonen S, Kononov A, Aittokallio T, Giri AK. SynToxProfiler: An interactive analysis of drug combination synergy, toxicity and efficacy. PLoS Comput Biol 2020; 16. doi:10.1371/JOURNAL.PCBI.1007604.

45 Reed LJ, Muench H. A simple method of estimating fifty per cent endpoints. Am J Epidemiol 1938; 27: 493–497.

46 Bolger AM, Lohse M, Usadel B. Trimmomatic: a flexible trimmer for Illumina sequence data. Bioinformatics 2014; 30: 2114.

47 Langmead B, Salzberg SL. Fast gapped-read alignment with Bowtie 2. Nat Methods 2012; 9: 357.

48 Li H, Handsaker B, Wysoker A, Fennell T, Ruan J, Homer N et al. The Sequence Alignment/Map format and SAMtools. Bioinformatics 2009; 25: 2078–2079.

49 Grubaugh ND, Gangavarapu K, Quick J, Matteson NL, De Jesus JG, Main BJ et al. An amplicon-based sequencing framework for accurately measuring intrahost virus diversity using PrimalSeq and iVar. Genome Biol 2019; 20: 1–19.

50 Paschoalino M,Marinho M dos S, Santos IA, Grosche VR, Martins DOS, Rosa RB, et al. An update on the development of antiviral against Mayaro virus: from molecules to potential viral targets. Archives of Microbiology 2023 205:4 2023; 205: 1–18.

51 Ogorek TJ, Golden JE. Advances in the Development of Small Molecule Antivirals against Equine Encephalitic Viruses. Viruses 2023; 15. doi:10.3390/V15020413.

52 Battisti V, Urban E, Langer T. Antivirals against the Chikungunya Virus. Viruses 2021; 13: 1307.

53 Hucke FIL, Bugert JJ. Current and Promising Antivirals Against Chikungunya Virus. Front Public Health 2020; 8: 618624.

54 Kovacikova K, van Hemert MJ. Small-Molecule Inhibitors of Chikungunya Virus: Mechanisms of Action and Antiviral Drug Resistance. Antimicrob Agents Chemother 2020; 64. doi:10.1128/AAC.01788-20.

55 Langendries L, Abdelnabi R, Neyts J, Delang L. Repurposing drugs for mayaro virus: Identification of eidd-1931, favipiravir and suramin as mayaro virus inhibitors. Microorganisms 2021; 9. doi:10.3390/MICROORGANISMS9040734/S1.

56 Ehteshami M, Tao S, Zandi K, Hsiao HM, Jiang Y, Hammond E et al. Characterization of β-d-N4-Hydroxycytidine as a Novel Inhibitor of Chikungunya Virus. Antimicrob Agents Chemother 2017; 61. doi:10.1128/AAC.02395-16.

57 Rosales-Rosas AL, Soto A, Wang L, Mols R, Fontaine A, Sanon A et al. β-D-N4-hydroxycytidine (NHC, EIDD-1931) inhibits chikungunya virus replication in mosquito cells and ex vivo Aedes aegypti guts, but not when ingested during blood-feeding. Antiviral Res 2024; 225: 105858.

58 Urakova N, Kuznetsova V, Crossman DK, Sokratian A, Guthrie DB, Kolykhalov AA et al. β-d-N4-Hydroxycytidine Is a Potent Anti-alphavirus Compound That Induces a High Level of Mutations in the Viral Genome. J Virol 2018; 92. doi:10.1128/JVI.01965-17.

59 Painter GR, Bowen RA, Bluemling GR, DeBergh J, Edpuganti V, Gruddanti PR et al. The prophylactic and therapeutic activity of a broadly active ribonucleoside analog in a murine model of intranasal venezuelan equine encephalitis virus infection. Antiviral Res 2019; 171: 104597.

60 Delang L, Abdelnabi R, Neyts J. Favipiravir as a potential countermeasure against neglected and emerging RNA viruses. Antiviral Res 2018; 153: 85–94.

61 Ferreira AC, Reis PA, de Freitas CS, Sacramento CQ, Hoelz LVB, Bastos MM et al. Beyond Members of the Flaviviridae Family, Sofosbuvir Also Inhibits Chikungunya Virus Replication. Antimicrob Agents Chemother 2019; 63. doi:10.1128/AAC.01389-18.

62 Mumtaz N, Jimmerson LC, Bushman LR, Kiser JJ, Aron G, Reusken CBEM et al. Cell-line dependent antiviral activity of sofosbuvir against Zika virus. Antiviral Res 2017; 146: 161–163.

63 Wichit S, Diop F, Hamel R, Talignani L, Ferraris P, Cornelie S et al. Aedes Aegypti saliva enhances chikungunya virus replication in human skin fibroblasts via inhibition of the type I interferon signaling pathway. Infect Genet Evol 2017; 55: 68–70.

64 Growth of human hepatoma cells lines with differentiated functions in chemically defined medium - PubMed. https://pubmed.ncbi.nlm.nih.gov/6286115/ (accessed 9 Jan2025).

65 Kabinger F, Stiller C, Schmitzová J, Dienemann C, Kokic G, Hillen HS et al. Mechanism of molnupiravir-induced SARS-CoV-2 mutagenesis. Nat Struct Mol Biol 2021; 28: 740.

66 Abdelnabi R, Foo CS, Kaptein SJF, Zhang X, Do TND, Langendries L et al. The combined treatment of Molnupiravir and Favipiravir results in a potentiation of antiviral efficacy in a SARS-CoV-2 hamster infection model. EBioMedicine 2021; 72. doi:10.1016/J.EBIOM.2021.103595.

67 Fung A, Jin Z, Dyatkina N, Wang G, Beigelman L, Deval J. Efficiency of incorporation and chain termination determines the inhibition potency of 2’-modified nucleotide analogs against hepatitis C virus polymerase. Antimicrob Agents Chemother 2014; 58: 3636–3645.

68 Ramirez S, Li YP, Jensen SB, Pedersen J, Gottwein JM, Bukh J. Highly efficient infectious cell culture of three hepatitis C virus genotype 2b strains and sensitivity to lead protease, nonstructural protein 5A, and polymerase inhibitors. Hepatology 2014; 59: 395–407.

69 Sacramento CQ, De Melo GR, De Freitas CS, Rocha N, Hoelz LVB, Miranda M et al. The clinically approved antiviral drug sofosbuvir inhibits Zika virus replication. Sci Rep 2017; 7. doi:10.1038/SREP40920.

70 Zhou S, Long N, Rosenke K, Jarvis MA, Feldmann H, Swanstrom R. Combined Treatment of Severe Acute Respiratory Syndrome Coronavirus 2 Reduces Molnupiravir-Induced Mutagenicity and Prevents Selection for Nirmatrelvir/Ritonavir Resistance Mutations. J Infect Dis 2024; 230: 1380–1383.

71 Kabinger F, Stiller C, Schmitzová J, Dienemann C, Kokic G, Hillen HS et al. Mechanism of molnupiravir-induced SARS-CoV-2 mutagenesis. Nat Struct Mol Biol 2021; 28: 740–746.

72. Sanderson T, Hisner R, Donovan-Banfield I, Hartman H, Løchen A, Peacock TP et al. A molnupiravir-associated mutational signature in global SARS-CoV-2 genomes. Nature 2023 623:7987 2023; 623: 594–600.

73 Zhou S, Long N, Rosenke K, Jarvis MA, Feldmann H, Swanstrom R. Combined Treatment of Severe Acute Respiratory Syndrome Coronavirus 2 Reduces Molnupiravir-Induced Mutagenicity and Prevents Selection for Nirmatrelvir/Ritonavir Resistance Mutations. J Infect Dis 2024; 230. doi:10.1093/INFDIS/JIAE213.

