## Supplemental Material for "Combinations of approved oral nucleoside analogues confer potent suppression of alphaviruses *in vitro* and *in vivo*"

Figures S1-S8

Table S1-S2.

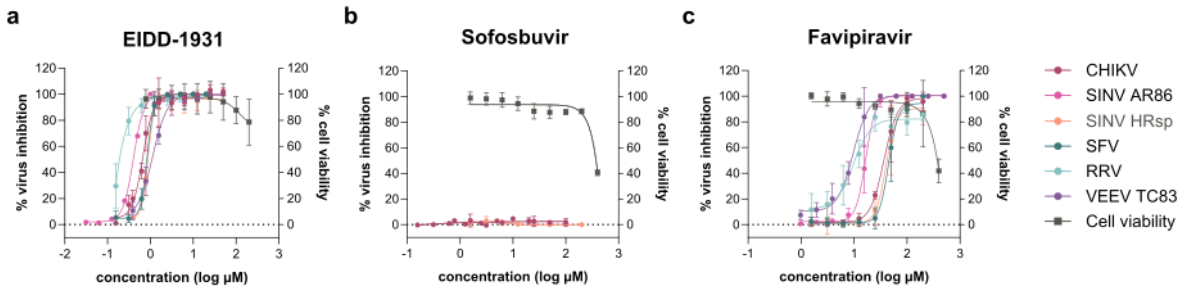


**Figure S1. Antiviral efficacy of single compounds against alphaviruses in Vero cells.** Dose-response activity of (a) EIDD-1931 (MPV), (b) SOF and (c) FAV against cytopathic effect induced by CHIKV, SINV, SFV, RRV or VEEV, quantified in Vero cells by the MTS method at 72 hours post infection. Data represent percentage of virus inhibition or cell viability compared to virus control and cell control, respectively, and is shown as mean values ± standard deviation from at least three independent experiments.

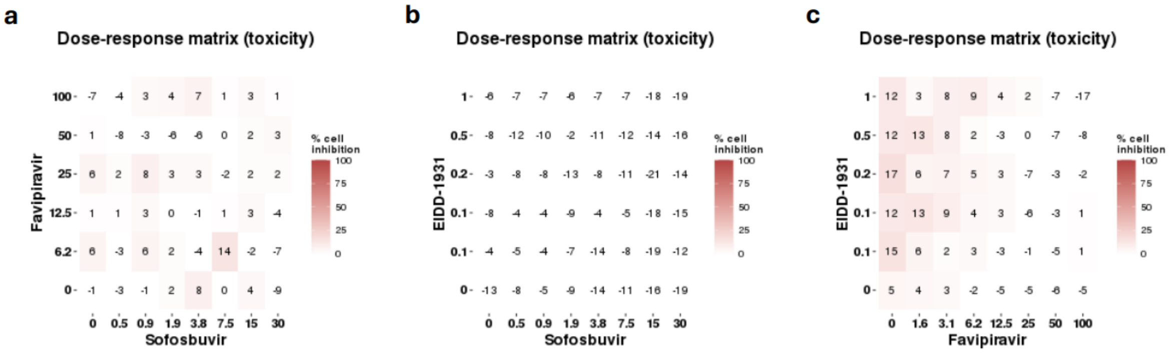


**Figure S2. Cytotoxicity of drug combinations in human skin fibroblasts.** Dose-response effect of combinations of (a) FAV and SOF, (b) MPV (EIDD-1931) and SOF, or (c) MPV and FAV on human skin fibroblasts, quantified by the MTS method at 72 hours post infection. Data present percentage cell death compared to cell control, analyzed using SynToxProfiler.

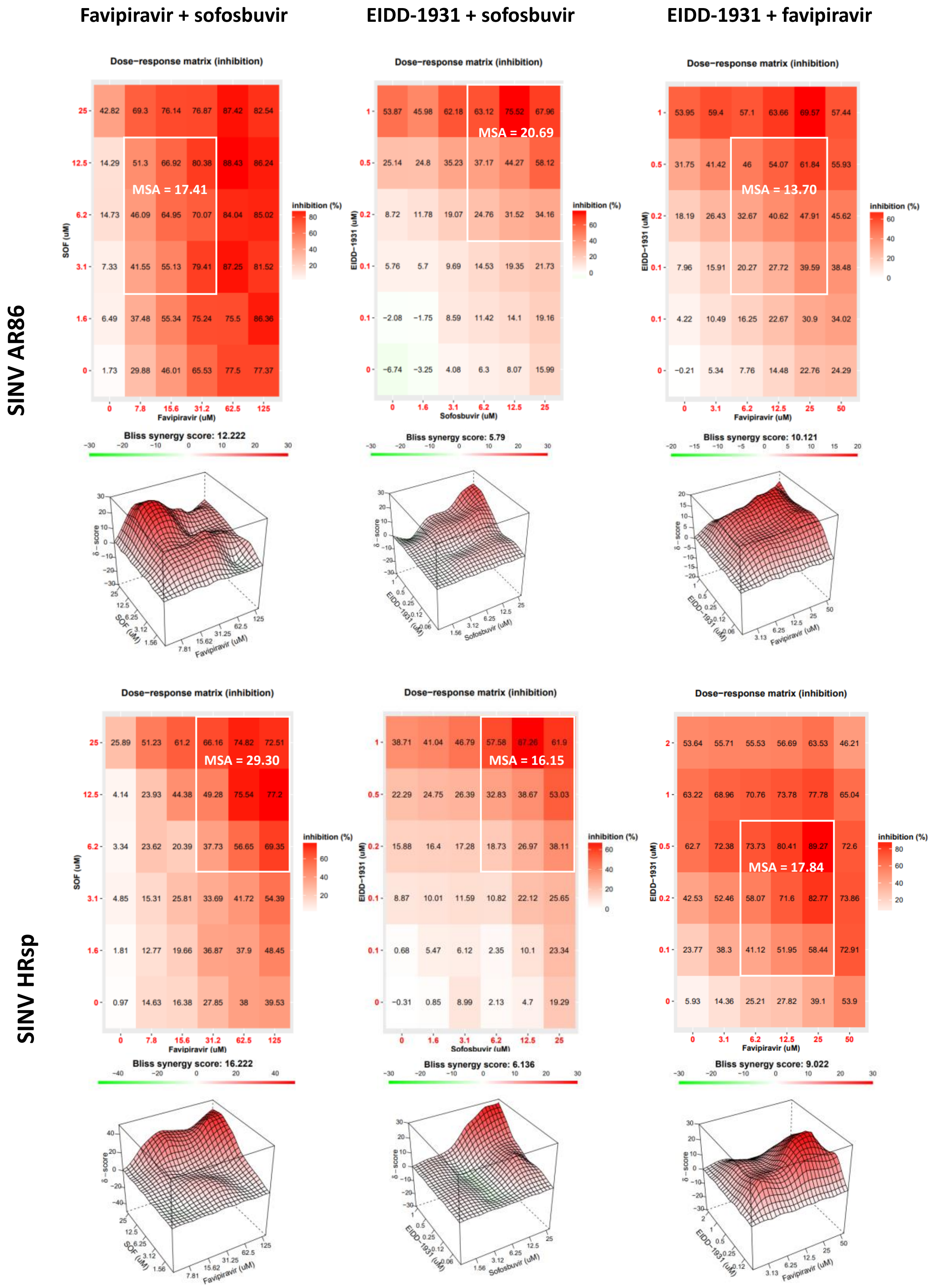


**Figure S3. Antiviral efficacy of combination treatment with DAAs against SINV in Huh7 cells.** Antiviral efficacy of combinations of EIDD-1931 (MPV) and FAV, EIDD-1931 (MPV) and SOF or FAV and SOF against SINV AR86 and HRsp strains in Huh7 cells. Inhibition of virus-induced CPE was quantified by the ATP method at 72 hours post infection. Data were analyzed and visualized using SynergyFinder based on the Bliss independence model. Dose-response matrices show percentage virus inhibition for each combination. MSA is indicated within white squares. Three-dimensional maps highlight the areas of synergy (red) and antagonism (green) across the full dose response matrix for each combination. Data comprise representative SF3.0 plots from single experiments. MSA, most synergistic area.

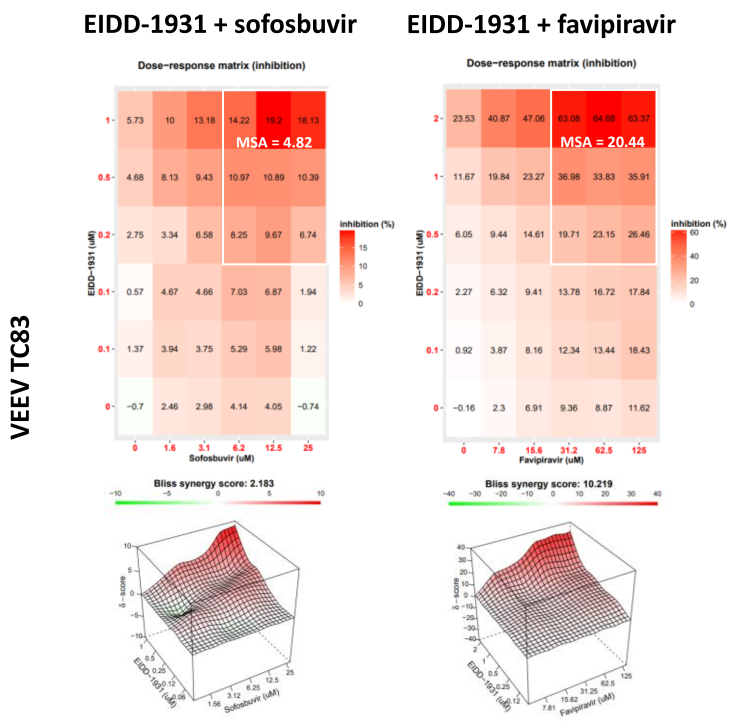


**Figure S4. Antiviral efficacy of combination treatment with DAAs against VEEV in Huh7 cells.** Antiviral efficacy of combinations of EIDD-1931 (MPV) and FAV, EIDD-1931 (MPV) and SOF or FAV and SOF against VEEV-TC83 in Huh7 cells. Inhibition of virus-induced CPE was quantified by the ATP method at 72 hours post infection. Data were analyzed and visualized using SynergyFinder based on the Bliss independence model. Dose-response matrices show percentage virus inhibition for each combination. MSA is indicated within white squares. Three-dimensional maps highlight the areas of synergy (red) and antagonism (green) across the full dose response matrix for each combination. Data comprise representative SF3.0 plots from single experiments. MSA, most synergistic area.

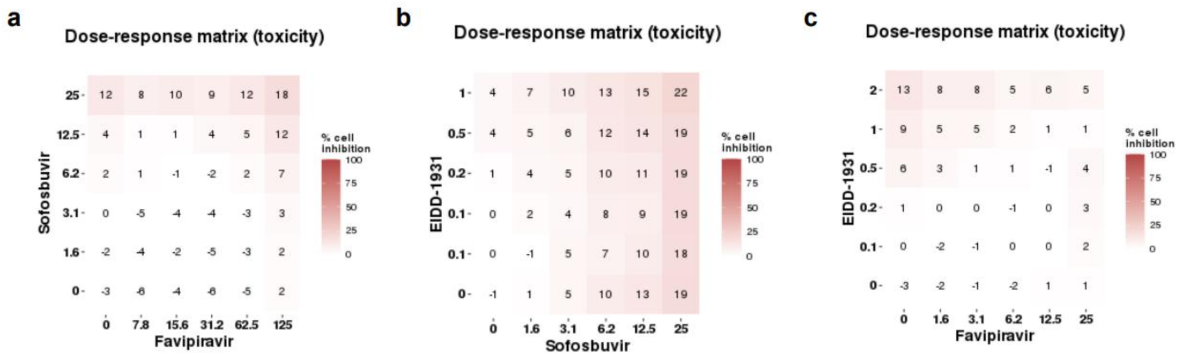


**Figure S5. Cytotoxicity of combination treatment with DAAs in Huh7 cells.** Dose-response effect of combinations of (a) FAV and SOF, (b) MPV (EIDD-1931) and SOF, or (c) MPV (EIDD-1931) and FAV on human skin fibroblasts, quantified by the ATP method at 72 hours post infection. Data present percentage cell death compared to cell control, analyzed using SynToxProfiler.

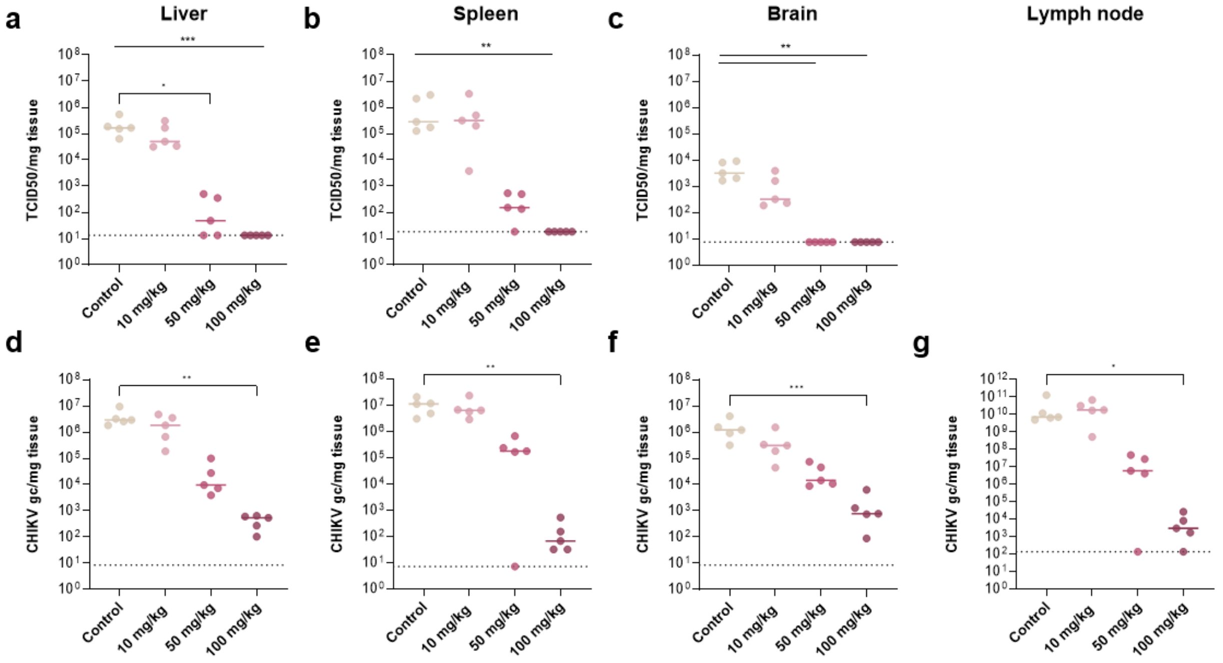


**Figure S6. Dose-dependent effect of MPV on CHIKV loads in AG129 mice.** AG129 mice (n=5 per group) were treated orally with different single doses of MPV (10, 50, 100 mg/kg) and infected subcutaneously with CHIKV (100 PFU) in the left hind foot. (a-c) Infectious virus titers in tissues on day 3 pi, quantified by end-point titrations on Vero cells. (d-g) Viral RNA levels in tissues on day 3 pi, quantified by qRT-PCR. Virus loads in the (a,d) liver, (b,e) spleen, (c,f) brain, and (g) draining lymph node on day 3 pi. Individual data points are shown, with solid lines representing median values. Statistical significance was assessed with Kruskal-Wallis test with Dunn's correction (*, $p<0.05$; **, $p<0.01$; ***, $p<0.005$). (a-c) Dotted lines represent the LOQ; (d-g) dotted lines represent the LOD. Gc, genome copies; TCID50, tissue culture infectious dose 50; pi, post infection; LOQ, limit of quantification; LOD, limit of detection.

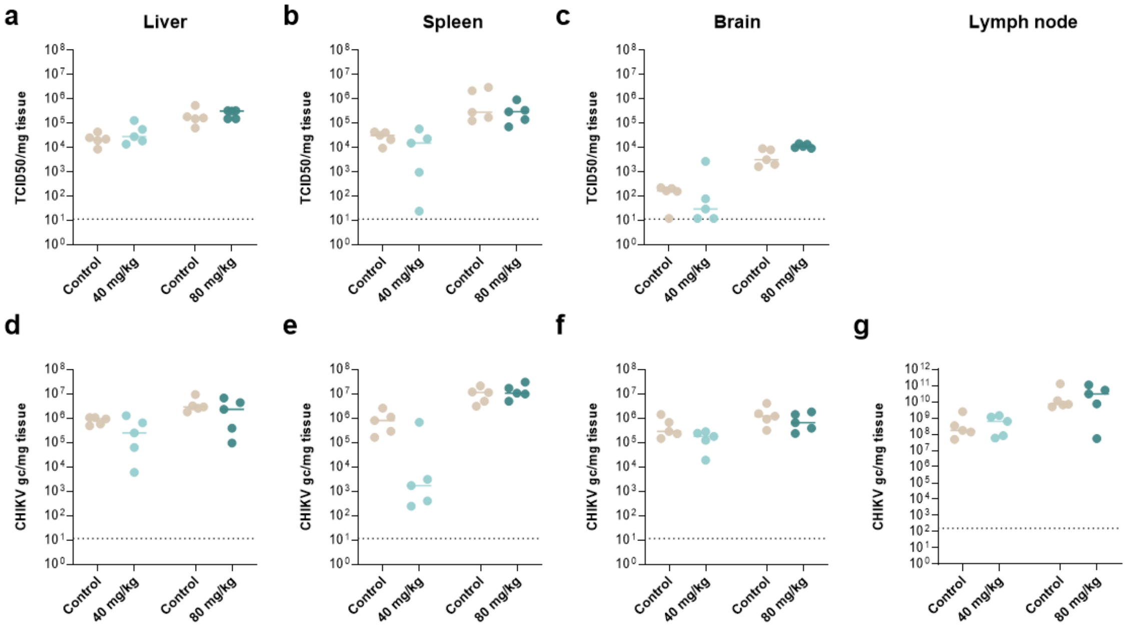


**Figure S7. Dose-dependent effect of SOF on CHIKV loads in AG129 mice.** AG129 mice (n=5 per group) were treated orally with different single doses of SOF (40, 80 mg/kg) and infected subcutaneously with CHIKV (100 PFU) in the left hind foot. (a-c) Infectious virus titers in tissues on day 3 pi, quantified by end-point titrations on Vero cells. (d-g) Viral RNA levels in tissues on day 3 pi, quantified by qRT-PCR. Virus loads in the (a,d) liver, (b,e) spleen, (c,f) brain, and (g) draining lymph node on day 3 pi. Individual data points are shown, with solid lines representing median values. Statistical significance was assessed with Kruskal-Wallis test with Dunn's correction (ns, $p>0.05$). (a-c) Dotted lines represent the LOQ; (d-g) dotted lines represent the LOD. Gc, genome copies; TCID50, tissue culture infectious dose 50; pi, post infection; LOQ, limit of quantification; LOD, limit of detection.

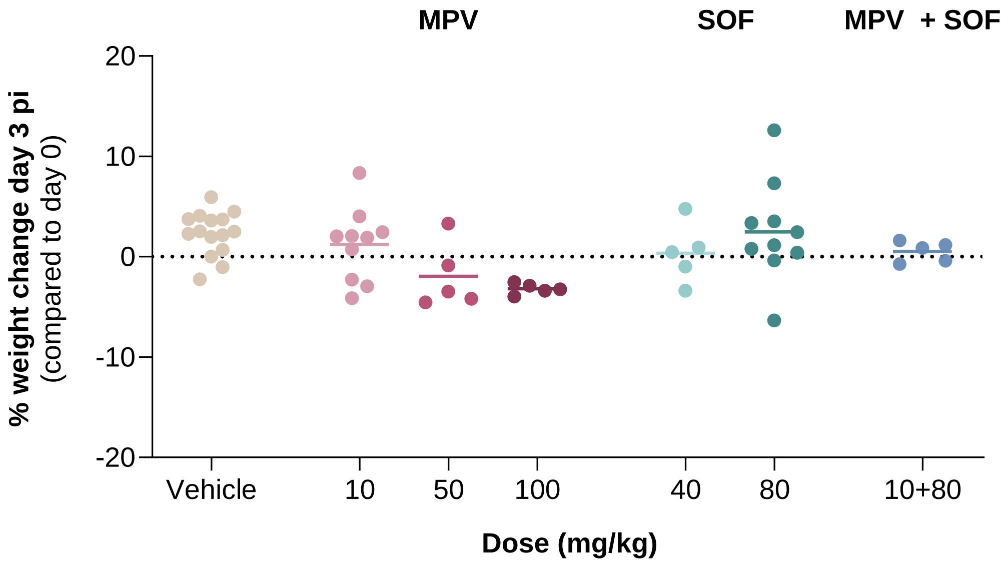


**Figure S8. Effect of single and combination treatments on body weight of AG129 mice.** AG129 mice (n=5 per group) were treated orally with single or combined doses of MPV and SOF and infected subcutaneously with CHIKV (100 PFU) in the left hind foot. Body weight was monitored daily and weight change at day 3 pi was calculated in percentage and normalized to the body weight at the time of infection. Kruskal-Wallis test with Dunn's correction was used to assess statistical significance (ns, $p>0.05$).

| Combination | Virus | Cell line | Overall Bliss Synergy score | MSA |
|---|---|---|---|---|
| MPV + SOF | CHIKV | Skin fibroblasts | 10.5 | 25.7 |
| FAV + SOF | CHIKV | Skin fibroblasts | 14.3 | 33.1 |
| MPV + FAV | CHIKV | Skin fibroblasts | 3.7 | 15.5 |
| MPV + SOF | SFV | Skin fibroblasts | 2.3 | 16.4 |
| FAV + SOF | SFV | Skin fibroblasts | 14.2 | 36.8 |
| MPV + FAV | SFV | Skin fibroblasts | 7.5 | 22.6 |
| MPV + SOF | VEEV-TC83 | Huh7 | 1.8 | 3.9 |
| MPV + FAV | VEEV-TC83 | Huh7 | 10.3 | 22.5 |
| MPV + SOF | SINV AR86 | Huh7 | 7.9 | 14.6 |
| FAV + SOF | SINV AR86 | Huh7 | 9.3 | 14.9 |
| MPV + FAV | SINV AR86 | Huh7 | 9.4 | 18.0 |
| MPV + SOF | SINV HRsp | Huh7 | 6.7 | 13.5 |
| FAV + SOF | SINV HRsp | Huh7 | 14.3 | 26.4 |
| MPV + FAV | SINV HRsp | Huh7 | 9.0 | 14.4 |

**Table S1.** Overview of Bliss Synergy scores and MSAs, from n=2-3 independent experiments. MSA; most synergistic area.

| Mutation | Group 1 | Group 2 | n1 | n2 | statistic | p | p.adj | p.adj.signif |
|---|---|---|---|---|---|---|---|---|
| T to C | COMBO | MPV10 | 5 | 4 | 5 | 0.13 | 0.13 | ns |
| T to C | COMBO | MPV50 | 5 | 5 | 0.5 | 0.008 | 0.023 | * |
| T to C | COMBO | MPV100 | 5 | 3 | 0 | 0.017 | 0.025 | * |
| G to A | COMBO | MPV10 | 5 | 4 | 7 | 0.265 | 0.265 | ns |
| G to A | COMBO | MPV50 | 5 | 5 | 0 | 0.006 | 0.017 | * |
| G to A | COMBO | MPV100 | 5 | 3 | 0 | 0.017 | 0.025 | * |
| C to T | COMBO | MPV10 | 5 | 4 | 9.5 | 0.5 | 0.5 | ns |
| C to T | COMBO | MPV50 | 5 | 5 | 1 | 0.008 | 0.024 | * |
| C to T | COMBO | MPV100 | 5 | 3 | 3 | 0.125 | 0.188 | ns |
| A to G | COMBO | MPV10 | 5 | 4 | 5.5 | 0.162 | 0.162 | ns |
| A to G | COMBO | MPV50 | 5 | 5 | 5 | 0.071 | 0.106 | ns |
| A to G | COMBO | MPV100 | 5 | 3 | 0 | 0.018 | 0.054 | ns |

**Table S2.** Results of Wilcoxon test on mutations induced in CHIKV RNA upon MPV treatment.
